# FRG1 Regulates Nonsense-Mediated mRNA Decay by Modulating UPF1 Levels

**DOI:** 10.1101/2025.11.18.689056

**Authors:** Ananya Palo, Talina Mohapatra, Anamika Singh, Shithij Thalakkat, Suryasikha Mohanty, Rajeeb Kumar Swain, Manjusha Dixit

**Affiliations:** National Institute of Science Education and Research, School of Biological Sciences, Bhubaneswar, Odisha 752050, India; Homi Bhabha National Institute, Training School Complex, Anushakti Nagar, Mumbai 400094, India; Institute of Life Sciences, Nalco Square, Chandrasekharpur, Bhubaneswar, Odisha 751023, India

**Author notes:** **Corresponding author:** Manjusha Dixit, PhD Associate Professor Room No. 204, School of Biological Sciences, National Institute of Science Education and Research, Bhubaneswar PO: Jatani, Khurda 752050, Odisha, India Phone: Office 91-674 2494195, Lab 91-674-2494196 Email ID.

**Keywords:** FSHD region gene 1, NMD, DUX4, UPF1, Structural component, Proteasomal degradation

## Abstract

FRG1 acts as a molecular double-edged sword: while its overexpression is linked to facioscapulohumeral muscular dystrophy (FSHD), it’s under expression activates tumorigenic signalling pathways—highlighting the importance of tightly regulated FRG1 levels. Although prior studies have hinted at FRG1’s involvement in RNA biogenesis, its core molecular functions have remained largely elusive. Our previous work identified FRG1 as a transcriptional regulator of genes in the nonsense-mediated decay (NMD) pathway; however, the underlying mechanisms were not fully delineated.

In this study, using a range of cellular models with altered FRG1 expression, we demonstrate that reduced FRG1 levels enhance NMD activity. We reveal that FRG1 is a structural component of both the spliceosome and the exon junction complex (EJC), and that it directly modulates NMD by interacting with UPF1—regulating its ubiquitination and degradation. Furthermore, we show that DUX4 inversely regulates the NMD machinery through FRG1, establishing a critical molecular axis. Despite its structural association with the EJC and spliceosome, the absence of FRG1 does not compromise the integrity of these complexes. Polysome profiling showed that FRG1 co-sediments with eIF4A3, a core EJC component, across translating ribosomal fractions. This interaction was further supported by proximity ligation assays confirming the close spatial proximity of FRG1 and eIF4A3. Importantly, we validated the impact of FRG1 perturbation on NMD efficiency and UPF1 levels in vivo using a transgenic FRG1 knockout zebrafish model.

Together, these findings establish a direct, DUX4-independent role for FRG1 in regulating the NMD pathway and uncover a previously unrecognized function for FRG1 in post-transcriptional gene regulation. Our work provides mechanistic insight into FRG1’s molecular roles and lays the groundwork for therapeutic strategies targeting diseases associated with its dysregulation.

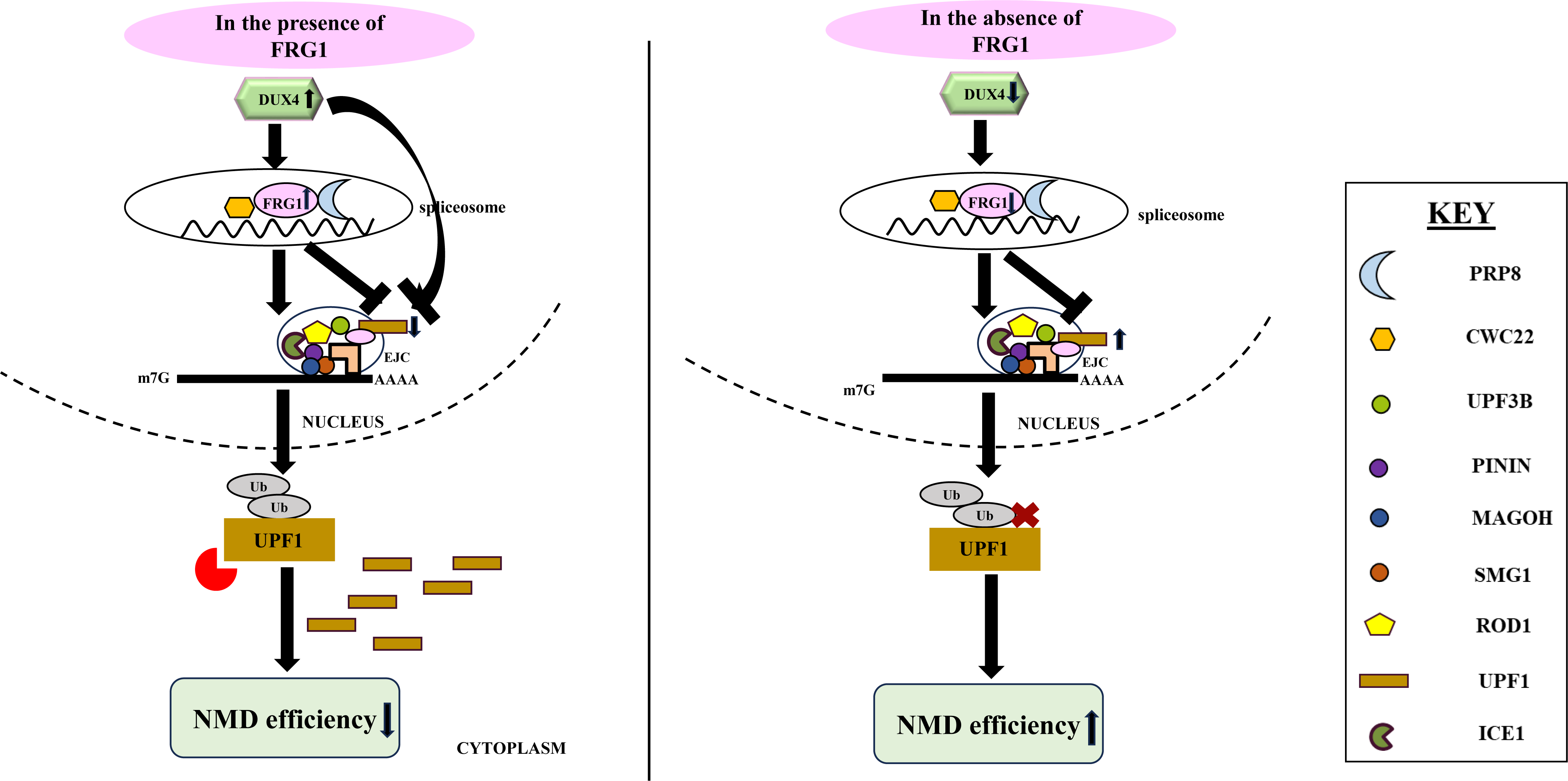
Graphical Abstract.

## Introduction

In eukaryotic organisms, gene expression undergoes quality control at various stages. One pivotal mechanism of RNA quality control is the Nonsense-mediated mRNA decay (NMD) pathway. NMD plays a critical role in preserving cellular equilibrium by degrading messenger RNAs (mRNAs) containing premature termination codons (PTCs), thus preventing the synthesis of potentially harmful proteins. Interest in NMD has been further fuelled by the studies showing the regulation of half-life and, thereby steady-state abundance of nearly 5-10% of mammalian transcriptome having upstream open reading frame (uORFs) in the 5’UTR, presence of long 3’UTR or 3’UTR harboring an intron, presence of uORF that interferes with the expression of main ORF, and downstream EJCs [1,2]. NMD is known to regulate various biological processes, including development, stress response, cell differentiation, and cell proliferation [3–9]. The loss of key NMD factors-UPF1, UPF2, or SMG1-results in embryonic lethality in mice, highlighting their essential role [10–12]. Evidence also suggests differential NMD activity across cell and tissue types. For instance, analysis of NMD reporters and various NMD substrates demonstrated higher NMD efficiency in HeLa cells compared to MCF7 cells [13]. Furthermore, RNAi-mediated depletion of NMD factors such as UPF1 or SMG1 led to DNA damage and inhibited the growth of HeLa cells, but not fibroblasts [14]. Together, these findings indicate that NMD is a highly regulated and context-dependent process.

The importance of NMD pathway has been highlighted as numerous genetic diseases that result from misexpression of faulty mRNAs escaping the mRNA surveillance pathway. Mutations in NMD factor UPF3B result in neurodevelopmental disorder, ADHD, autism, and schizophrenia [15–17]. Dysregulation of NMD factors like UPF2, UPF3A, SMG6, eIF4A3, RNPS1 results in intellectual disability (ID) disorder [18,19]. Aberrant NMD is associated with cancer [20,21]. For example, mutations in UPF1 lead to inflammatory myofibroblastic tumors (IMT) and pancreatic cancer [22,23]. Deletion of one allele of SMG1 results in lung cancer and hematopoietic malignancies [24]. Upregulation of AKT results in increased cancer cell proliferation [25–27]. Additionally, tumors may benefit from NMD-mediated degradation by acquiring PTCs in tumor suppressor genes like WT1, TP53, RB, and BRCA1/2 [28–30].

This dual role of NMD in degrading aberrant mRNAs and post transcriptionally targeting a subset of normal mRNAs is believed to occur through different pathways. Gehring et al. identified two branches of NMD - UPF2 and RNPS1 or EJC dependent [31]. Subsequently, Chen et al. identified a branch of NMD pathway that is independent of UPF3B [32]. Ultimately, the SURF complex assembles on the aberrant transcripts which are degraded via the DECID complex [33–36]. Despite significant progress in understanding the NMD pathway, critical gaps remain, particularly in deciphering the precise molecular mechanisms that govern its specificity and efficacy. The emerging evidence of a broader role for NMD in gene expression regulation raises pressing questions: How is the NMD response buffered? How are transcripts without PTCs selectively targeted by the NMD machinery? And, importantly, is the NMD pathway itself subject to regulatory control? Addressing these gaps is essential for a comprehensive understanding of NMD’s role in cellular processes and disease. In the past, with an intent to discover novel NMD genes that may aid in therapeutic interventions for genetic diseases, primary genetic screens were conducted in model organisms that resulted in identification of factors such as ROD1, ICE1, DHX34, NBAS [37–39]. Immunoprecipitation to identify the interacting proteins of NMD factor ROD1 (EJC component) unveiled the binding of FRG1 with the former [37], hinting towards a direct or indirect role played by FRG1 (FSHD Region Gene 1) in NMD.

FRG1 is a gene with increased expression leading to facioscapulohumeral muscular dystrophy (FSHD) [40–42] while reduced expression is linked to several types of cancer including oral, gastric, colorectal, prostrate, and breast cancer [43–46]. Some reports suggest potential role of FRG1 in RNA biogenesis. FRG1 has been found to localise within the nucleolus in HeLa cells and associates with Cajal bodies, nucleoli and nascent transcripts in *Xenopus laevis* oocytes [47]. RIP-based assay in HeLa and C2C12 cells shows the direct binding of FRG1 with FXR1 and Rbfox1 mRNA, respectively [47,48]. Yeast two hybrid assays have shown FRG1 to be a part of spliceosome Bact/ C complex [49]. In the same study, co-immunoprecipitation revealed binding of CWC22 with FRG1 and eIF4A3. CWC22 is an essential splicing factor that initiates EJC assembly via direct interaction with EJC core protein eIF4A3 [50]. Cryo EM structure by Bertram et al. showed intricate spatial organisation of FRG1 within spliceosome C complex. Here, FRG1 was shown to interact with NTD (N-terminal domain) of splicing factor PRP8 [51]. The aforementioned studies showing FRG1’s interaction with spliceosome and EJC gave a plausible hint of its role in the post transcriptional processing of transcripts. Additionally, our previous work revealed that FRG1 depletion affects transcript level of NMD factors (ICE1, PNN, MAGOH, ROD1, UPF3B, SMG1) having roles in different steps of the NMD pathway [52]. However, whether FRG1 affects NMD, and if so, whether FRG1 acts as a structural component of the NMD machinery, remains unclear, as does the exact mechanism by which FRG1 regulates the NMD pathway within the cell.

In this manuscript, we have employed NMD loss-of-function and gain-of-signal luciferase reporter systems, as well as an intron-retention splicing construct, to investigate the impact of altered FRG1 expression on nonsense-mediated mRNA decay (NMD) activity. We have utilized both FRG1 knockout and overexpression cellular models to systematically examine its role in modulating NMD efficiency. Additionally, we have performed transcriptome analyses to assess changes in the expression of known NMD target transcripts, particularly those harboring premature termination codons (PTCs). To probe the structural and functional associations of FRG1 within the spliceosome–EJC–NMD machinery, we conducted co-immunoprecipitation, polysome profiling, and proximity ligation assays. Furthermore, we evaluated the in vivo relevance of our in vitro experiments using a transgenic frg1 knockout zebrafish model. Our findings offer new insights confirming FRG1’s role in modulating the NMD pathway, and its underlying molecular mechanism.

## Materials and Methods

### Zebrafish maintenance

AB strain of zebrafish was used in this study. The animal work was approved by the Institutional Animal Ethics Committee. Inclusion criteria were fertilized one-cell stage embryos from healthy adult breeders (6–12 months; female:male ratio 2:1) with normal morphology and survival beyond 24 hpf. Exclusion criteria included unfertilized eggs, developmental arrest, abnormal morphology, or infection. Embryos were raised in E3 media (34.8 gm NaCl, 1.6 gm KCl, 5.8 gm CaCl2.2H2O, 9.78 gm MgCl2.6 H2O; pH-7.2) at 28°C. At 24 hpf, embryos were bleached using 0.001% sodium hypochlorite solution (Sigma Aldrich, MO, USA) to prevent any kind of infection. These were then transferred into a circulating water system tank (Techniplast, Odisha, India) at 5 dpf. Embryos were randomized by clutch, with microinjections performed by one investigator and phenotyping assessed by blinded evaluators.

### frg1 knockout generation

To generate frg1 knockout transgenic zebrafish line, sgRNAs targeting frg1 exon1 and exon 9 (sequences mentioned in Supplementary Table S1) were designed following the protocol by Varsheney et al. [53]. The gene-specific oligos were annealed with the 5’-AAAAGCACCGACTCGGTGCCACTTTTTCAAGTTGATAACGGACTAGCCTTATTTTAACTTGCTATTTCTAGCTCTAAAAC-3’ (gRNA backbone) using Phusion polymerase (Thermo Scientific, IL, USA). The guide RNAs were in vitro transcribed using MEGAscript™ SP6 Transcription Kit (Invitrogen) and Megascript^TM^ T7 transcription kit (Invitrogen, MA, USA), respectively. Cas9 mRNA was synthesized by linearizing the pCs2-nCas9n nanos 3’ UTR vector (Addgene, #62542) with XbaI-HF (NEB, MA, USA) followed by in vitro transcription using mMESSAGE mMACHINE^TM^ transcription kit (Invitrogen). The adult male and female fishes were bred, and eggs were collected at 1-celled stage. A Femtojet microinjector (Eppendorf, Hamburg, Germany) was used to inject embryos with a mixture containing 25 ng/µl each of exon1 and exon 9 sgRNAs, along with 100 ng/µl Cas9 mRNA.

### Screening of frg1 knockout zebrafish embryos

Embryos were genotyped three months post-injection to screen for mutation incorporation, using caudal fin clipping across F0, F1, and F2 generations. Genomic DNA was isolated using 1x TE buffer (10 mM Tris-HCl, pH 7.5, and 1 mM EDTA, pH 8.0) along with proteinase K treatment (Roche, Basel, Switzerland), followed by PCR using exon-1 forward and exon-9 reverse primers with primers listed in Supplementary Table S1.

### Whole mount in-situ hybridisation (WISH)

WISH was performed on zebrafish embryos as described earlier (Thisse and Thisse, 2008) [54]. Briefly, the zebrafish frg1 coding sequence (Supplementary Table S1) was amplified and cloned in pCR™-Blunt II-TOPO (Invitrogen). The DIG (Digoxigenin) labelled RNA probe was synthesized by linearizing the plasmid with BamH1 (NEB) and transcribing with T7 polymerase. The embryos were then fixed, dehydrated through methanol series (25% to 100%) and stored at -20°C in 100% methanol.-After rehydration, the embryos were incubated in a hybridization buffer containing 50% deionized formamide, 5X SSC, 50 µg/ml heparin, 0.5 mg/ml torula RNA, 9.2 mM citric acid, and 0.1% Tween-20, along with a zebrafish FRG1-specific probe. The following day, embryos were incubated in 2% blocking reagent (Roche-1109617600) for 2 hours at room temperature, to prevent non-specific binding. Staining of embryos was done by incubating them with anti-digoxigenin (Roche) FAB fragment coupled with alkaline phosphate diluted in blocking buffer. On the 3^rd^ day, embryos were washed with MABT (Maleic Acid buffer with Tween) and pH9 buffer at room temperature and then transferred into BM purple dye (Roche) for color development, in dark.

### Cell culture, Transfection and stable line generation

MCF7 cells were obtained from Cell Repository, NCCS, Pune, India. The cell line was authenticated by the repository using 16 STR markers (AmpFISTR Identifier Plus PCR Amplification Kit, Applied BioSystems). We checked mycoplasma contamination using MycoAlert Mycoplasma Assay kit (Lonza). The cell line was grown in DMEM media (Himedia, Mumbai, India) supplemented with 10% FBS (Himedia, US origin) and 1X Penicillium-Streptomycin-Amphotericin B (PSA; Himedia, Mumbai, India). Cells were cultured at 37°C and 5% CO_2_. FRG1 knockdown (pLKO.1_FRG1sh, TRCN0000075012, RRID: AB_259684) and expression (HsCD004 21091 PLX304_FRG1) vectors were procured from Sigma and Harvard repository, respectively. Transfection of the cells was done with the constructs using Lipofectamine 3000 according to the manufacturer’s protocol (Invitrogen).

Single-cell colonies were selected for overexpression and knockdown using blasticidin (10 µg/ml) and puromycin (2 µg/ml) (Sigma Aldrich, MO, USA) antibiotics, respectively. Confirmation of the stable lines was done using Western Blotting.

### NMD assay

NMD sensitive luciferase reporter (pCMV-3XFLAG-Fluc-betaglobin-39PTC) was purchased from Addgene repository (Addgene:#112084). pKC-4.06 plasmid was a kind gift from Dr. James Inglese’s lab (pKC-4.06 plasmid was a kind gift from Dr. James Inglese’s lab RRID: Addgene_112084) [38]. The NMD sensor plasmid was microinjected into 1-cell stage zebrafish embryos. It was also transfected in MCF7 perturbed cells (MCF7_FRG1_overexpressed, MCF7_FRG1_Knockout and MCF7_FRG1_Knockdown) and their corresponding control cells (MCF7_Empty Vector, MCF7_Wild type and MCF7_Scrambled), respectively using Lipofectamine 3000 (Invitrogen, MA, USA) in a 12 well plate at 50% cell confluency. 48 h post transfection, cell lysate was prepared according to manufacturer’s protocol (Promega, WI, USA). Luciferase signal was recorded using ^®^Varioska^®^ Flash Multimode Reader (Thermo Scientific, MA, USA). Experiment was performed in triplicate.

### Co Immunoprecipitation

MCF7 cells (11 million) were plated and grown in 150 mm dishes. The cells were washed with DPBS (Himedia) and fixed using 1% formaldehyde (Himedia) for 10 minutes at room temperature. Reaction was stopped using 1.25M glycine solution (MP Biomedicals, OH, USA). Cells were scrapped after adding ice-cold PBS and lysed using non-denaturing mammalian lysis buffer (20 mM Tris, pH-7.5, 150 mM NaCl, 1 mM EDTA, 1 mM EGTA, 1% NP-40) along with protease inhibitor (Invitrogen). Cell lysate was prepared by sonicating cells at 40% amplitude, 30 sec OFF/ON cycle, 5 times using a Ultrasonic Processor Sonicator (Cole-Parmer, IL, USA). The cell lysate was centrifuged at maximum speed for 10 mins at 4°C to collect the supernatant. The supernatant was divided into three equal parts-whole cell lysate, IgG (negative control), and test sample (antibody of interest). Immunoprecipitation was performed overnight at 4°C by incubating the samples with 5 µg of respective primary antibodies [(FRG1, Abcam, MA, USA), (GAPDH, Abgenex, India), (DUX4, Novus Biologicals, CO, USA), (UPF1, CST, MA, USA), eIF4A3 (Abcam, MA, USA), CWC22 (Abcam, MA, USA), IgG (H+L) (Rockland, USA), PRP8 (Abcam, MA, USA), UPF3B (Abcam, MA, USA)]. Following day, protein A agarose bead (Santa Cruz Biotechnology, TX, USA) was washed with 1x PBS and incubated with the samples at 4°C for 8 hours. Post incubation, beads were extensively washed with ice cold PBS, boiled for 5 min with Laemlli buffer. Subsequently, immunoblotting was carried out. The bands were detected by chemiluminescence using SuperSignal^TM^ West Femto maximum sensitivity substrate (Thermo Scientific). Band intensities were normalised to GAPDH.

### Proteasome inhibition

In a 12-well plate, 0.1 million MCF7 cells having DUX4 overexpression were plated in three wells. Post 12 hours, FRG1 expression vector (1 µg) was transfected in one of the wells using Lipofectamine 3000 (Invitrogen). Induction with 500 ng/ml doxycycline (MP Biomedicals) was done for a period of 24 hours. Post 16 hours of induction, MG132 (50 µM) (Sigma Aldrich, MA, USA) was added in this condition (DUX4 induction along with FRG1 expression). Subsequently, western blotting was carried out to visualise the proteasomal inhibition.

### Western Blotting

Whole cell protein lysate was prepared using ice-cold RIPA buffer (Thermo Scientific, IL USA) and a cocktail of protease and phosphatase inhibitor (Thermo Scientific). BCA protein assay kit (Thermo Scientific) was used to quantify the total protein concentration. 30 µg of protein samples were loaded and resolved on 12% sodium dodecyl sulfate polyacrylamide gel electrophoresis (SDS-PAGE) and transferred onto PVDF membrane (Millipore, Bangalore, India). Membrane blocking was done using 5% skimmed milk (MP Biomedicals) for an hour and incubated overnight with respective antibodies of interest. Next day, blot washing was done using 1x TBST buffer and incubated with HRP conjugated secondary antibody for one hour. Blots were again washed with 1x TBST buffer and detected in Chemidoc using SuperSignal^TM^ West Femto maximum sensitivity substrate (Thermo Scientific). Images were analysed in Image J software (NIH, MD, USA) RRID: SCR_003070. All experiments were performed in triplicate.

### Quantitative Real Time PCR

Total RNA was isolated from MCF7 cells. frg1 transgenic knockout and AB strain wild type zebrafish using RNeasy mini kit (Qiagen, Hilden, Germany). Reverse transcription was performed using verso cDNA synthesis kit (Thermo Scientific, Lithuana) with 1µg of total RNA according to manufacturer’s instructions. Each RT-PCR reaction was carried out using 20ng cDNA, 2x SYBR Green PCR Master Mix (Thermo Scientific, CA, USA), and respective primers (Supplementary Table S1) using QS-7 system (Applied Biosystem-Thermo Scientific, CA, USA). The experiment was carried out in triplicate using GAPDH as an internal control (cell lines) and EF1A (zebrafish). Fold change was calculated using 2^-ΔΔCT^ method.

### Library preparation for RNA-seq

Total RNA was isolated from MCF7 FRG1 perturbed cell lines (knockout and knockdown) following manufacturer’s instructions as described above. RNA was quantified using NanoDrop One^c^ 2000 (Thermo Scientific). Purity of the samples were assessed using 2100 Bioanalyzer (Agilent Technologies, Waldbronn, Germany). Samples with RNA Integrity Number (RIN>8) were considered for library preparation. Sequencing libraries were prepared using 1 μg of total RNA for each sample with the Illumina TruSeq Stranded Total RNA Library Prep Human/Mouse/Rat Kit (Illumina, CA, USA). mRNA was isolated from these samples using magnetic beads with mRNA isolation kit (Poly A mRNA isolation module, NEB). Next, this mRNA was used to synthesize double-stranded cDNA using NEBNext Ultra II RNA library Preparation kit, according to the manufacturer’s instructions. Then indexing adapters were ligated to each sample. To generate NGS libraries, the products were enriched using PCR with adapter universal primers and purified using SPRIselect beads. The prepared libraries were quantified and checked for fragment size using the Qubit High Sensitivity DNA Reagent (Qubit 2.0) followed by TapeStation D1000 ScreenTape (Agilent Technologies). Sequencing of the resulting NGS libraries were performed using the NextSeq550 platform (llumina).

### Transcript Quantification, Isoform ratio measurements, and PTC prediction

The quality of the RNA-Seq data was assessed using FastQC (v0.12.2, RRID: SCR_014583)) to ensure high-quality sequencing reads. Transcript quantification was performed using Salmon (version 1.7.0) for all the samples by using the Salmon generated *decoy-aware* transcriptome index prepared using the Gencode SCR_014966 reference files (Release 47; GRCh38/hg38) reference files. The command included parameters specifying the library type: ISF, enabling validation of mappings, and correcting for sequence-specific biases [55]. The expressed isoform from each gene, as determined by the transcripts per million (TPM) from Salmon, was used for downstream analyses.

To investigate isoform usage and identify transcripts containing PTCs, quantified RNA-Seq data were analyzed using IsoformSwitchAnalyzeR (version 2.2.0), an R package [56]. Isoforms were categorized based on annotations from GENCODE release 47. IsoformSwitchAnalyzeR identifies NMD-sensitive transcripts using the 50-nucleotides rule [57–60].

Significant isoform switches were determined by applying thresholds on the differential isoform fraction (|dIF| > 0.1) and FDR corrected P-value (q-value; q < 0.05). Q-values were computed using the DEXSeq-based statistical framework implemented in IsoformSwitchAnalyzeR [61,62]. To facilitate visualization, illustrating isoform usage changes between conditions, density plots were generated using the ggplot2 (version 3.5.1, RRID-SCR_014601) package in R. Data set overlaps were analyzed using the ComplexHeatmap (version 2.18.0) package of R and visualized as an UpSet plot [63]. All scripts for the RNA-Seq analysis and NMD-prediction are available at GitHub https://github.com/sugar-syrup/FRG1-NMD.

### Polysome Assay

MCF7 cells were maintained in complete medium at 37 °C and treated with 100 μg/mL cycloheximide for 30 min to stabilize polysomes. Cells were washed with ice-cold PBS containing cycloheximide, lysed in hypotonic buffer (200 mM Tris-HCl [pH 7.4], 100 mM KCl, 5 mM MgCl₂, 1% Nonidet P-40, 5 mM DTT, 40 U/mL RNasin, and protease inhibitors), and layered onto 10–40% sucrose gradients (50 mM Tris-HCl [pH 7.4], 5 mM MgCl₂, 100 mM KCl). Lysates were ultracentrifuged at 36,000 rpm for 2 h at 4 °C using Beckman SW41 rotor. Gradients were fractionated using a fractionator (Biocomp gradient station and fraction collector) with continuous monitoring at 254 nm. Proteins from each fraction were ethanol-precipitated, resuspended in SDS sample buffer, resolved by 12% SDS-PAGE, and analyzed via immunoblotting.

### Proximity Ligation Assay

To assess protein–protein interactions, MCF7 cells were fixed in 4% paraformaldehyde and permeabilized with 0.2% Triton X-100. ( RRID:CVCL_0031) Following permeabilization, cells were blocked using 2% bovine serum albumin (BSA) in PBS to reduce nonspecific binding. Cells were then incubated with primary antibodies targeting FRG1 (mouse monoclonal, Abnova, Taiwan) and eIF4A3 (rabbit monoclonal, Abcam, USA), raised in different host species to enable dual recognition. The proximity ligation assay was performed using the Duolink® In Situ PLA Red Kit (Cat. No. DUO92101, Sigma-Aldrich) according to the manufacturer’s instructions. Species-specific PLA probes (PLUS and MINUS) were applied, and if the target proteins were in close proximity (<40 nm), the probes ligated to form a circular DNA molecule. This DNA was subsequently amplified by rolling circle amplification and visualized as distinct red fluorescent puncta under a fluorescence microscope. The number of dots indicates the level of protein–protein interaction. As a negative control, parallel samples were processed without the addition of primary antibodies. These control samples underwent the same procedure with PLA probes and amplification reagents to assess background signal due to nonspecific probe interaction or amplification.

See Supplementary Tables for detailed information on sources of reagents, software, and antibodies.

## Results

### FRG1 expression perturbation alters NMD efficacy

Based on our previous publication showing FRG1 as a direct transcriptional activator of NMD genes [52], we assessed the effect of its expression levels on NMD efficacy. To achieve this, we transiently transfected a specialized NMD sensor (pKC-4.06) in MCF7 cells having knockdown, knockout, and ectopic expression of FRG1. An increase in luciferase activity suggests impaired NMD machinery if PTCs are retained within transcripts in NMD sensor, and vice versa. As shown in Fig. 1 A-B, depletion and knockout of FRG1 resulted in decreased luciferase signal (0.8 and 0.83-folds, respectively) from the NMD sensor, indicating an increase in NMD efficiency. Ectopic expression of FRG1 resulted in increased luciferase signal (3.63-fold) from the NMD sensor, indicating reduced NMD efficiency (Fig. 1 C). To validate our *in vitro* observation, evaluating FRG1 expression perturbation on NMD efficacy, we generated FRG1 knockout zebrafish model. For this, we used CRISPR-Cas9 method, injecting sgRNAs targeting exon-1 and 3’UTR regions of frg1 in AB strain zebrafish (Fig. 1 D-G). This was designed to delete most of the entire frg1 locus in zebrafish. The knockout of frg1 in zebrafish was validated using PCR and Whole-mount in situ hybridization (Suppl. Fig. 1). To carry out NMD assay, 1-cell stage embryos were collected and microinjected with NMD sensor construct. Reporter activity between knockout and control zebrafish embryos showed enhanced NMD efficiency 48hour post fertilization (lane-2) (Fig. 1 H). Microinjecting the NMD sensor construct along with FRG1 overexpression plasmid in the frg1^-/-^ zebrafish embryos at 1-cell stage resulted in reduced NMD efficacy as observed at 48 hours post-fertilization (lane-3) (Fig. 1 H). These findings suggest that NMD is affected by FRG1.

**Figure 1.**
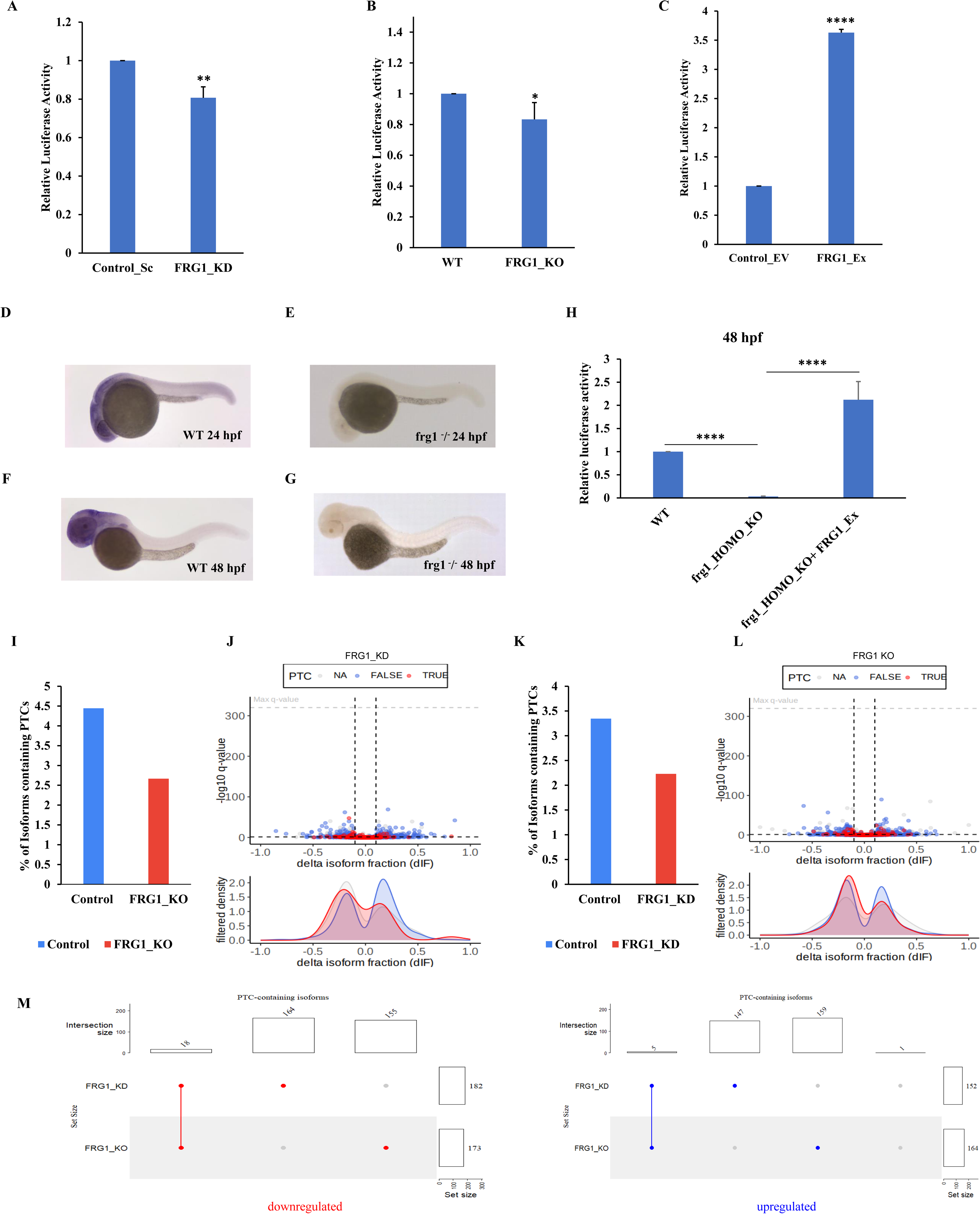
FRG1 depletion increases NMD efficiency. **(A–C)** Bar graphs depict relative luciferase activity in the NMD reporter under varying conditions of FRG1 manipulation: knockdown **(A)**, knockout **(B)**, and overexpression **(C)**. The figure uses the following abbreviations for labeling: Control_Sc (Control_Scrambled), Control_EV (Control_Empty Vector), WT (Wild Type), FRG1_KD (FRG1 Knockdown), FRG1_KO (FRG1 Knockout), and FRG1_Ex (FRG1 Overexpression). **(D-G)** Expression of frg1 in wild-type (WT) and frg1 knockout zebrafish at different time points. Image **(D)** and **(E)** show frg1 expression in WT and frg1 knockout embryos, respectively, at 24 hours post-fertilization (hpf). Image **(F)** and **(G)** depict frg1 expression in WT and frg1 knockout embryos, respectively, at 48 hpf. **(H)** The bar graph illustrates relative luciferase activity from NMD reporter in the zebrafish embryos having different frg1 expression conditions [wild type (WT), homozygous frg1 knockout (frg1_HOMO_KO), and FRG1 overexpression on frg1 homozygous knockout background (frg1_HOMO_KO+Ex)] at 48 hpf. **(I, K)** Bar graph depict the proportion of transcripts containing PTCs in MCF7 cell line having FRG1 knockdown **(I)** and knockout (K). The analysis was conducted using IsoformSwitchAnalyzeR on RNA-Seq data from MCF7 cells following FRG1 depletion. (J, L) The panels show Volcano plots, illustrating the change in isoform fraction (dIF) plotted against the −log10 of the q-value, highlighting differential transcript usage in FRG1-depleted RNA-Seq data from MCF7 cells. Isoforms are classified based on GENCODE (release 47) annotations: those with PTCs are marked in red (TRUE), with regular stop codons in blue (FALSE), and isoforms lacking an annotated open reading frame in gray (NA). The lower panels present density plots showing the distribution of filtered isoforms relative to thresholds of |dIF| > 0.1 and q-value < 0.05. Q-values were calculated using IsoformSwitchAnalyzeR’s DEXSeq-based statistical framework. (M) The UpSet plot illustrates the overlap of upregulated and downregulated isoforms containing PTCs identified in the FRG1 knockdown (KD) and knockout (KO) RNA-Seq datasets. Experiments were performed in triplicate. Results are presented as mean ± SD. ns-non-significant; * represents p-value ≤ 0.05; ** represents p-value ≤ 0.01; **** represents p-value ≤ 0.0001.

### FRG1 depletion regulates abundance of PTC-containing transcripts

Genome-wide RNA sequencing analysis was performed on FRG1 knockout, knockdown, and their respective control cells to assess whether increased NMD efficiency upon FRG1 depletion results in a decreased abundance of RNA isoforms containing premature translation termination codons. Fig. 1 I, J shows that the knockdown of FRG1 led to a reduced abundance of transcript isoforms containing PTCs. Among the top 10 genes having altered isoforms with functional consequences (NMD status), 50% exhibited a significant reduction in the NMD-sensitive isoforms upon FRG1 depletion (Fig. 2 A, B; Suppl. Fig. 2). Whereas only one gene having NMD-sensitive isoform showed a significant increase, and the remaining either showed no change in levels or non-significant change.

**Figure 2.**
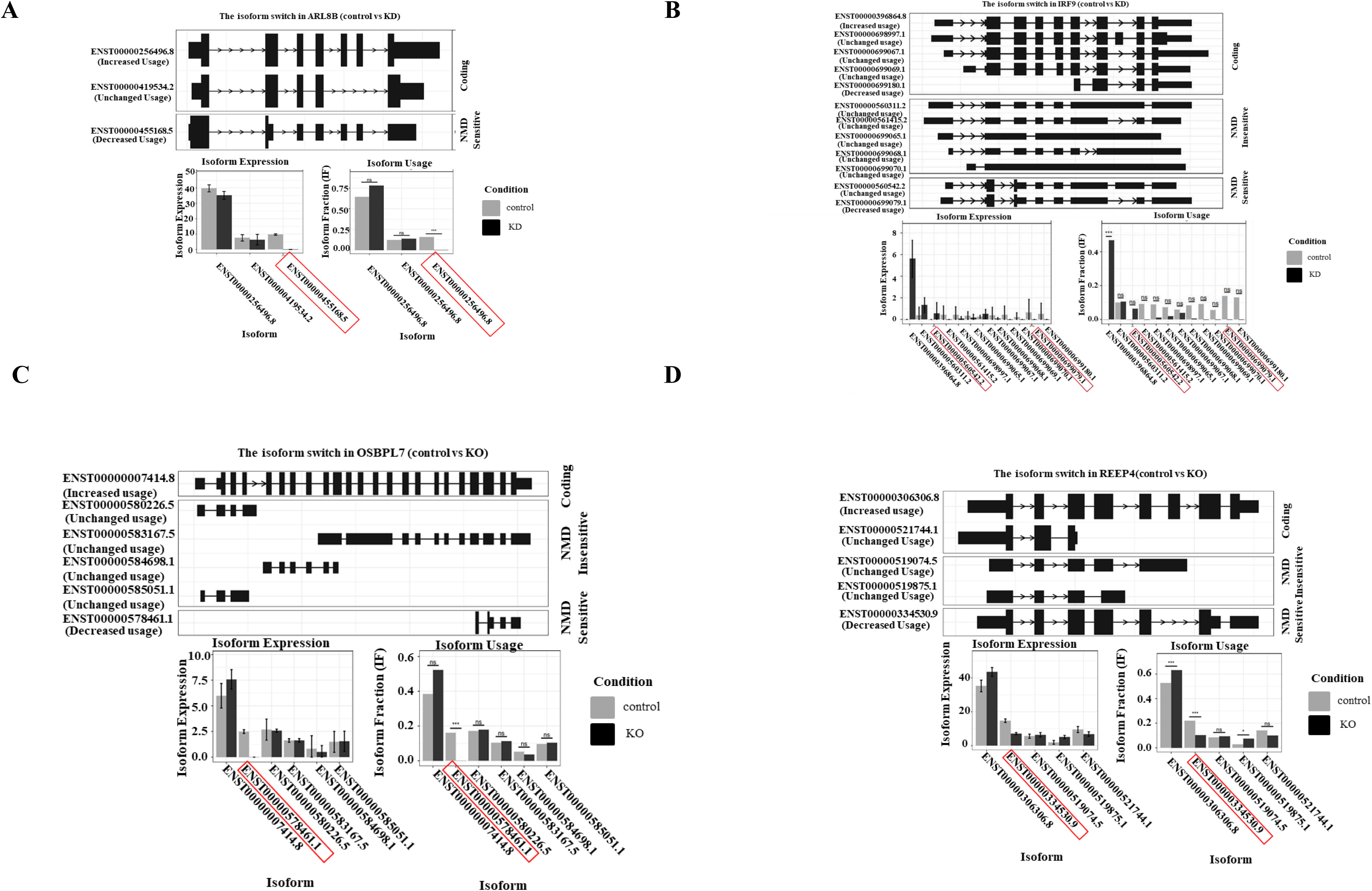
NMD-sensitive isoform levels upon FRG1 depletion. **(A-B)** The figure shows the isoform expression profile plots displaying isoform structures with annotated Ensembl IDs, isoform expression, and isoform fractions for genes ARL8B **(A)** and IRF9 **(B)**, in FRG1 knock down (**FRG1_KD**) set. **(C-D).** The figure shows the isoform expression profile plots illustrating the isoform structures with annotated Ensembl IDs, isoform expression levels, and isoform fractions for the genes OSBPL7 **(panel C)** and REEP4 **(panel D)** in the FRG1 knockout (FRG1_KO) dataset. Red boxes highlight the NMD-sensitive isoforms of the respective genes. (The remaining top 8 genes exhibiting isoform switching are shown in Suppl. Fig. 2 and 3 for FRG1 knockdown and knockout, respectively.)

The influence of FRG1 on NMD efficiency was notably more pronounced in its complete absence (Fig. 1 K, L). The knockout of FRG1 resulted in a substantial reduction in NMD-sensitive isoforms for 60% of the top 10 genes with functionally significant altered isoforms (NMD status). On the other hand, only two of these genes showed a significant increase in NMD-sensitive isoforms, while the others exhibited either a non-significant increase or no change (Fig. 2 C, D; Suppl. Fig. 3).

On a transcriptome-wide scale, comparing FRG1 depleted and knockout conditions, we found more overlap in NMD-sensitive isoforms showing downregulation (18) than upregulation (5) (Fig. 1 M). These findings provide evidence that FRG1 plays a crucial role in regulating the elimination of key NMD targets.

### DUX4 regulates NMD via FRG1

Feng et al. reported that DUX4, a transcriptional activator of FRG1, inhibits NMD [64–66]. Mechanistically, we investigated whether FRG1 directly influences NMD, potentially exceeding the impact of DUX4, or whether DUX4 exerts its effects on NMD independently of FRG1. To ascertain this, we created a stable, inducible MCF7 cell line that expresses DUX4 upon doxycycline induction. As reported previously [65], increased levels of DUX4 led to reduced NMD efficacy in MCF7 cells. Relative levels of luciferase signal obtained from the NMD sensor construct was 1.5-fold higher in the DUX4 overexpressing cells, showing the accumulation of PTC containing transcripts. However, NMD activity was restored (0.22-fold) with less intensity of luciferase signal from the NMD sensor construct due to decreased accumulation of PTC containing transcripts upon FRG1 depletion in DUX4 overexpressing cells (Fig. 3). This data suggested that the DUX4-mediated inhibition of NMD can be via FRG1.

**Figure 3.**
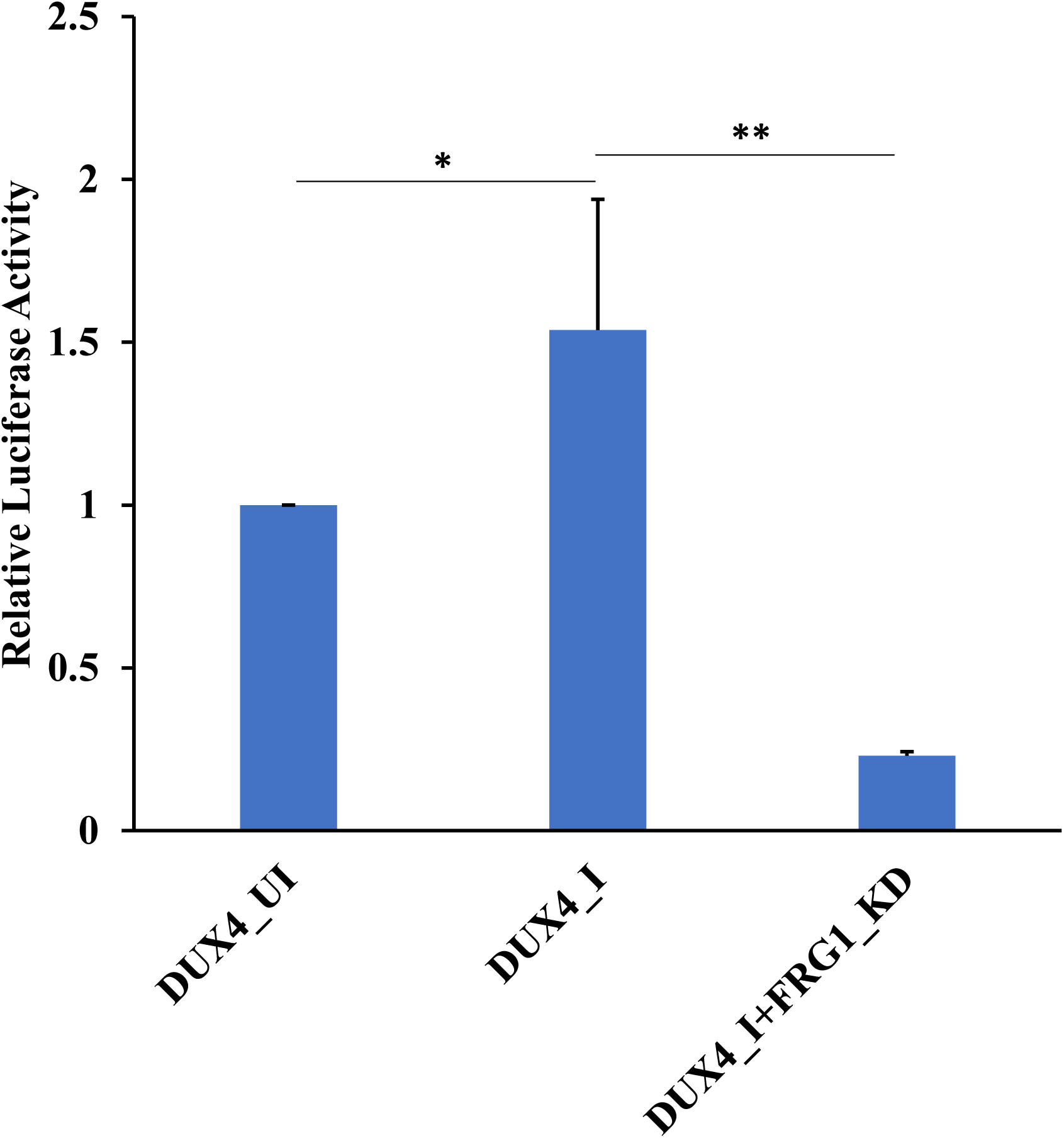
DUX4 modulates NMD efficiency via FRG1. The bar graph shows relative luciferase signal from NMD reporter plasmid under DUX4 and FRG1 perturbed conditions [DUX4 uninduced (DUX4_UI), DUX4 expression induced (DUX4_I), DUX4 expression induced and FRG1 knockdown (DUX4_I+FRG1_KD)] in MCF7 cells. Experiments were performed in triplicate. Results are presented as mean ± SD. * represents p-value ≤ 0.05; ** represents p-value ≤ 0.01.

### FRG1 dually regulates UPF1 expression, a central molecule in NMD

Since previous studies demonstrated that DUX4 influences UPF1 protein levels [65] and FRG1 affects the transcript levels of NMD factors [52], we investigated the impact of FRG1 on both UPF1 transcript and protein levels. We found an increase in UPF1 expression upon FRG1 reduction, both at mRNA (fold change = 1.93) (Fig. 4 A) and protein levels (Fig. 4 B). On the other hand, overexpression of FRG1 resulted in reduced UPF1 levels both at the RNA level (fold change =0.8) (Fig. 4 C) and protein level (Fig. 4 D). Parallel to *in vitro* data, knockout of frg1 in zebrafish resulted in increased upf1 expression (fold change=1.86) (Fig. 4 E). To ascertain the specificity, we ectopically expressed FRG1 in the frg1^KO/KO^ zebrafish model, which showed reduction in upf1 expression (fold change=0.36) (Fig. 4 F). This data indicated that FRG1 inversely regulates UPF1 expression.

**Figure 4.**
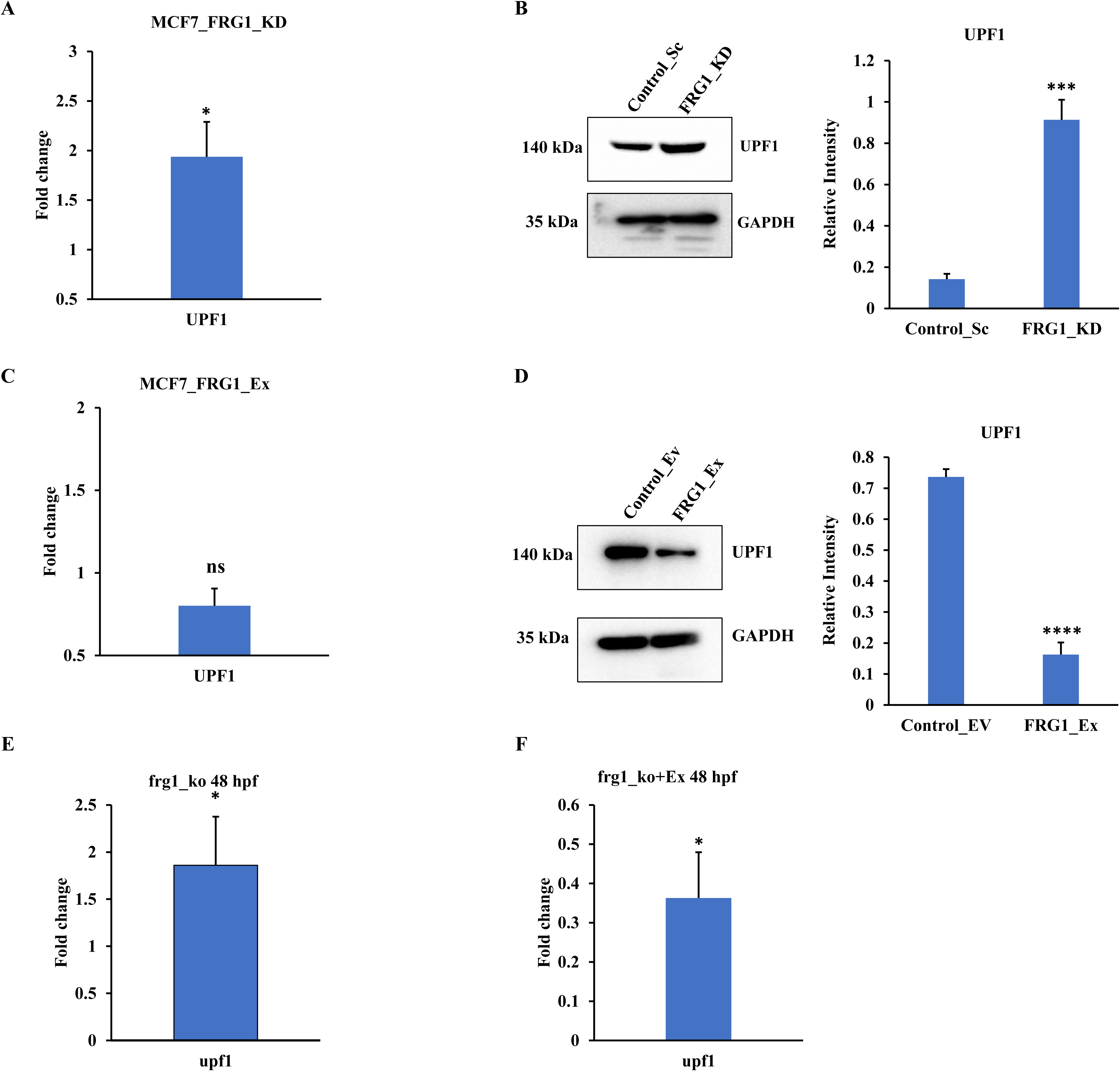
FRG1 expression perturbation alters UPF1 expression. **(A-B)** qRT-PCR expression data **(A)** and Western blot images **(B)** showing UPF1 expression upon FRG1 depletion in MCF7 cells. **(C-D)** qRT-PCR expression data **(C)** and Western blot images **(D)** showing UPF1 levels upon ectopic FRG1 expression in MCF7 cells. Here are the details of abbreviations used in labeling: FRG1 knockdown (FRG1_KD), FRG1 overexpression (FRG1_Ex), Control_Scrambled (Control_Sc)/ Control_Empty Vector (Control_EV). Experiments were performed in triplicate. **(E)** Bar graph depicts the qRT-PCR based expression levels of upf1 following knockout of frg1 (frg1_ko) in zebrafish. (F) Bar graph shows the qRT-PCR based expression levels of upf1 following ectopic expression of FRG1 under frg1 knockout background (frg1_ko+Ex) in zebrafish. Results are presented as mean ± SD. ns-nonsignificant; * represents p-value ≤ 0.05; *** represents p-value ≤ 0.001; **** represents p-value ≤ 0.0001.

### FRG1 Promotes Depletion of UPF1 Protein Independently of DUX4

Our previous publication demonstrated the direct binding of FRG1 to the promoter region of UPF1, establishing its role as a transcriptional regulator [52]. Building on these findings, we investigated whether FRG1 exerts a direct effect on UPF1 protein levels, a phenomenon previously reported in the context of DUX4. Consistent with earlier observations, DUX4 overexpression led to a marked reduction in UPF1 levels [65]. Interestingly, knockdown of FRG1 in a DUX4-overexpressing background restored UPF1 expression, indicating that FRG1 directly contributes to the downregulation of UPF1 protein levels (Fig. 5A). To further investigate this relationship, we overexpressed FRG1 in MCF7 cells, both with and without DUX4 overexpression. Notably, FRG1 overexpression alone led to a marked reduction in UPF1 levels—comparable to the reduction seen in cells already engineered to overexpress DUX4 (Fig. 5B).

**Figure 5.**
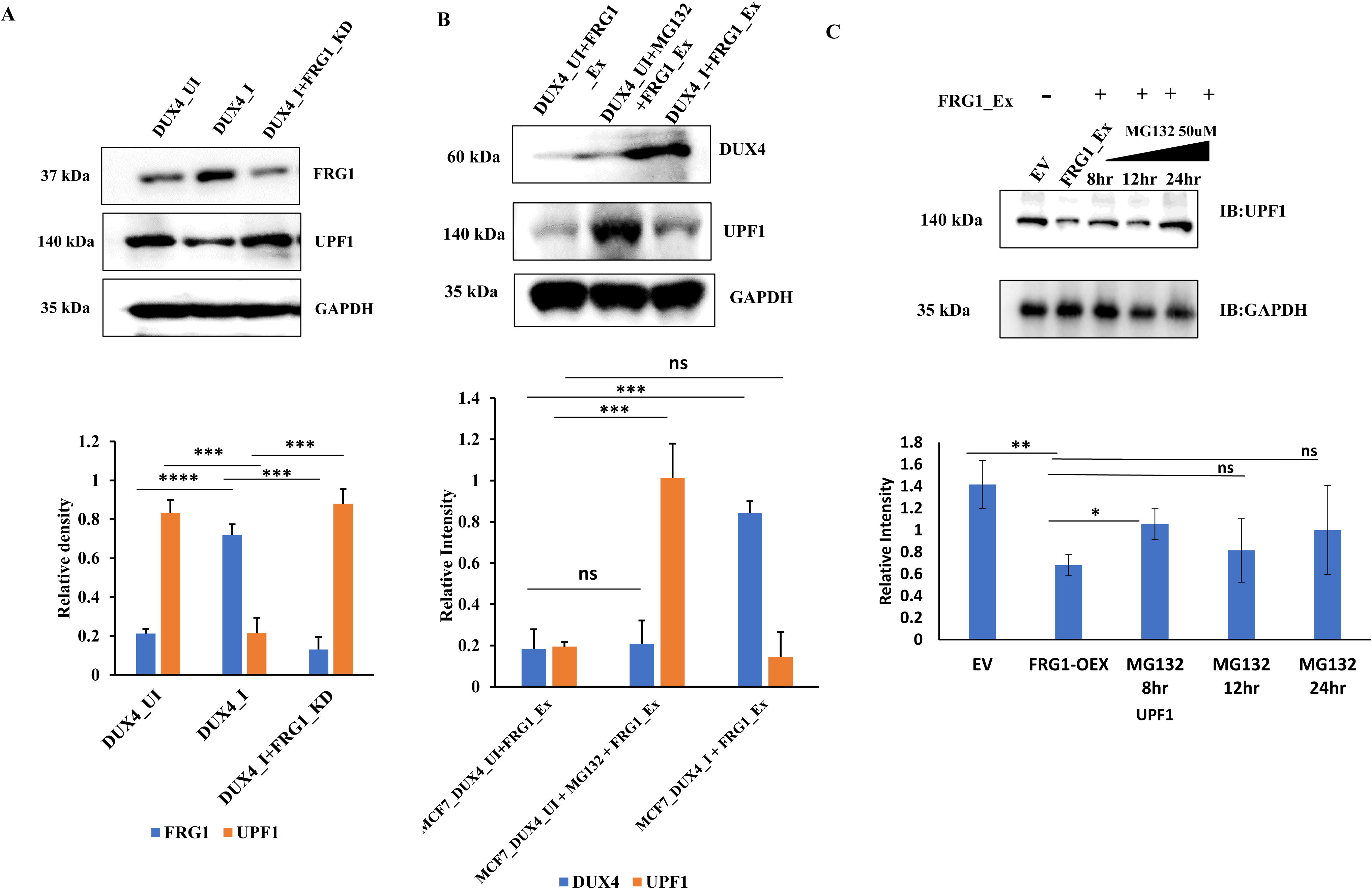
FRG1 Facilitates UPF1 Protein Reduction Independent of DUX4 Activity. **(A)** Western blot analysis illustrating UPF1 protein levels in MCF7 cells under various conditions: lane-2 shows the effect of DUX4 expression (DUX4_I), while subsequent lanes depict the impact of FRG1 knockdown in the presence of DUX4 expression (DUX4_I + FRG1_KD). **(B)** Western blot showing time-dependent effect of MG132 treatment (8, 12, and 24 h) on UPF1 ubiquitination in FRG1-overexpressing MCF7 cells. UPF1 was immunoprecipitated and probed with anti-ubiquitin antibody to assess polyubiquitination levels. (displays representative Western blot results showing UPF1 expression in MCF7 cells with endogenous DUX4 expression (DUX4_UI) and overexpression of FRG1 (FRG1_Ex), following treatment with MG132 (50 µM) (Lane-2). Lane-3 illustrates UPF1 expression in MCF7 cells co-expressing DUX4 and FRG1 (DUX4_I + FRG1_Ex). Experiments were performed in triplicate. Results are presented as mean ± SD. ns-nonsignificant; *** represents p-value ≤ 0.001; **** represents p-value ≤ 0.0001.C)

To explore the underlying mechanism, we examined the impact of proteasomal degradation. We found that inhibiting proteasomal degradation in FRG1-overexpressing cells restored UPF1 levels (Fig. 5B). Treatment of MCF7 cells overexpressing FRG1 with 50 µM MG132 for 8, 12, and 24 hours led to a time-dependent increase in UPF1 protein levels (Fig. 5C), confirming that UPF1 degradation is mediated via the proteasome.

Collectively, these results provide strong evidence that FRG1 directly promotes UPF1 protein depletion through a proteasome-dependent mechanism, independent of DUX4.

### FRG1 associates directly with UPF1 protein and promotes its ubiquitination and degradation

Next, we investigated whether FRG1 functions as part of the nonsense-mediated decay machinery and negatively regulates UPF1 levels through direct interaction. Co-immunoprecipitation assays, performed using reciprocal pulldown with either FRG1 or UPF1 antibodies, confirmed a protein-protein interaction between FRG1 and UPF1 (Fig. 6A–C). These findings suggest that FRG1 serves as a structural component of the NMD complex.

**Figure 6.**
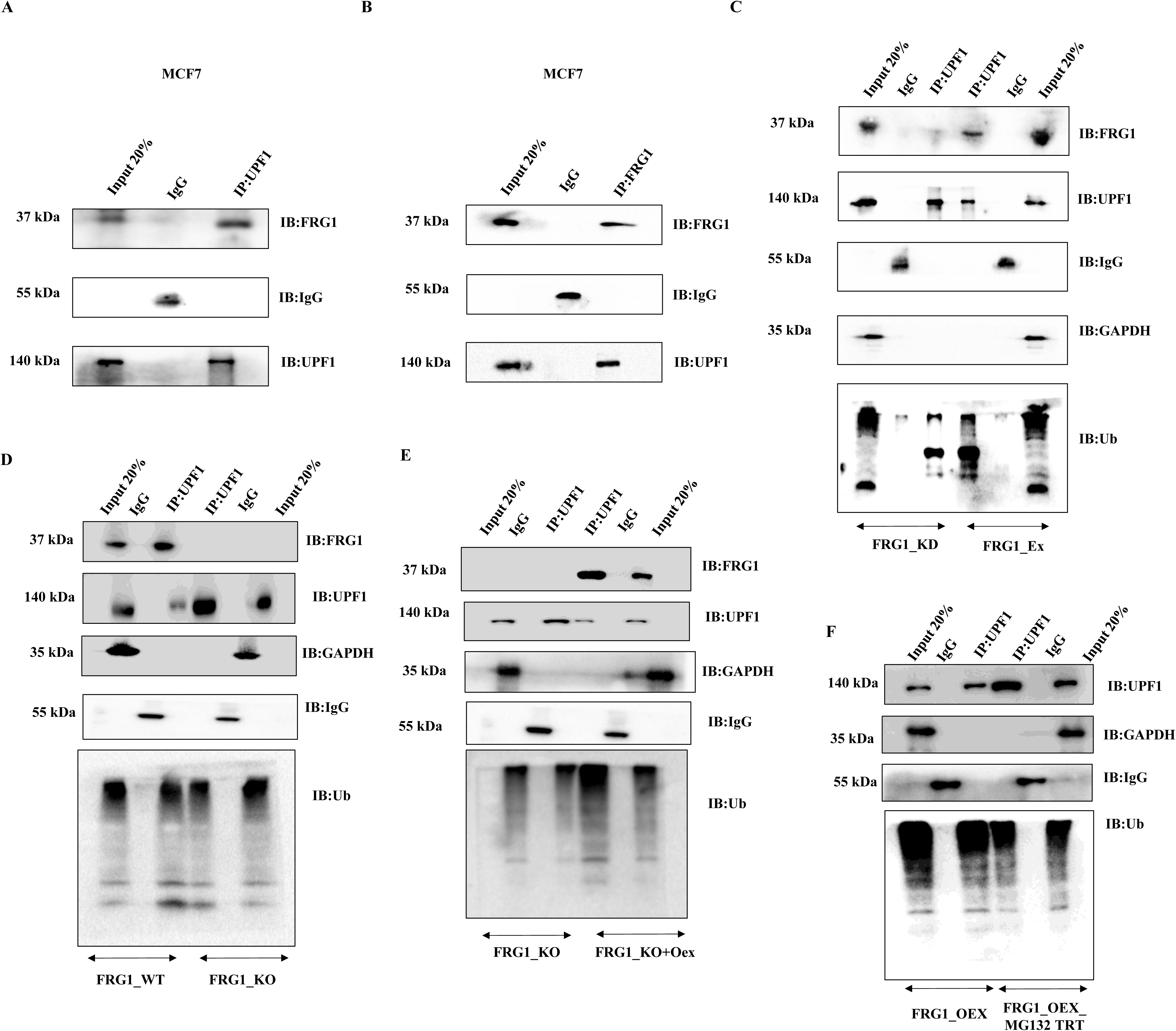
FRG1 interacts with UPF1 and mediates its proteasomal degradation. **(A-B)** Co-immunoprecipitation with UPF1 **(A)** and FRG1 **(B)** antibodies and representative Western blots showing interaction of FRG1 with UPF1 in MCF7 cell line. **(C)** Immunoblot analysis of a quantitative co-immunoprecipitation assay using UPF1 antibody demonstrates UPF1 ubiquitination under conditions of FRG1 knockdown (FRG1_KD, left three lanes) and FRG1 overexpression (FRG1_Ex, right three lanes). Co-immunoprecipitation with IgG was used as negative control. **(D-F)** Validation of FRG1-dependent regulation of UPF1 ubiquitination and stability via the proteasome pathway. **(D)** Immunoblot analysis of a quantitative co-immunoprecipitation assay using UPF1 antibody demonstrates UPF1 ubiquitination (UB) under conditions of FRG1 endogenous levels (FRG1_WT, left three lanes) and FRG1 knockout (FRG1_KO, right three lanes). Co-immunoprecipitation with IgG was used as negative control. **(E)** Immunoblot analysis of a quantitative co-immunoprecipitation assay using UPF1 antibody shows comparison of UPF1 ubiquitination (UB) in FRG1 knockout background (FRG1_KO left three lanes) with reintroduction of FRG1 in the knockout background (FRG1_KO+Oex, right three lanes) . **(F)** Immunoblot analysis of a quantitative co-immunoprecipitation assay using a UPF1 antibody compares UPF1 ubiquitination (UB) levels in FRG1-overexpressing cells (FRG1_Ex, left three lanes) with those in FRG1-overexpressing cells subjected to an 8-hour treatment with 50 µM MG132 (FRG1_OEX_MG132 TRT).

To further investigate the underlying mechanism, we performed a co-immunoprecipitation-based ubiquitination assay. FRG1 depletion led to a marked reduction in UPF1 ubiquitination, accompanied by increased binding between UPF1 and FRG1. In contrast, ectopic expression of FRG1 enhanced UPF1 ubiquitination while diminishing its interaction with FRG1 (Fig. 6D). To validate these findings, we employed FRG1 knockout MCF7 cells and observed that ectopic re-expression of FRG1 in this background successfully restored UPF1 ubiquitination, confirming the role of FRG1 in regulating this process (Fig. 6E). Furthermore, a UPF1 pulldown assay performed under conditions of FRG1 overexpression—following treatment with 50 µM MG132 for 8 hours—showed reduced levels of ubiquitinated UPF1 (Fig. 6F). This finding confirms a proteasome-mediated degradation pathway.

Collectively, these results demonstrate that FRG1 regulates UPF1 stability by promoting its ubiquitination and subsequent proteasomal degradation. Thus, FRG1 acts as a negative regulator of UPF1 levels via a ubiquitin–proteasome mechanism (Fig. 5G–I).

### FRG1 is a structural component of the EJC and spliceosome complex

FRG1 is reported as a structural component of spliceosomal C complex [49]. Overexpression of FRG1 leads to missplicing of transcripts [67]. Since splicing and nonsense-mediated mRNA decay are closely interconnected processes [68,69], and since components of the exon junction complex (EJC)—assembled during splicing—play a critical role in NMD [70], we investigated whether FRG1 influences NMD by altering UPF1 levels or by disrupting the structural integrity of the spliceosome and EJC. To explore this, we assessed the stability of these complexes in FRG1-perturbed cells. PRP8, a core catalytic component of the spliceosomal C complex [71], has been shown via cryo-electron microscopy (Cryo-EM) to interact with FRG1 [51]. CWC22, a pre-mRNA splicing factor, bridges splicing and the EJC [50]. We performed co-immunoprecipitation experiments on nuclear extracts from cells with altered FRG1 expression, using an antibody against PRP8. Immunoblotting confirmed that FRG1 interacts with both PRP8 and CWC22 in FRG1-overexpressing cells (Fig. 7 A). However, in FRG1-deficient cells, the interaction between PRP8 and CWC22 remained unaltered (Fig. 7 B). These findings suggest that although FRG1 is a structural component of the human spliceosomal C complex, its absence does not compromise the complex’s structural integrity.

**Figure 7.**
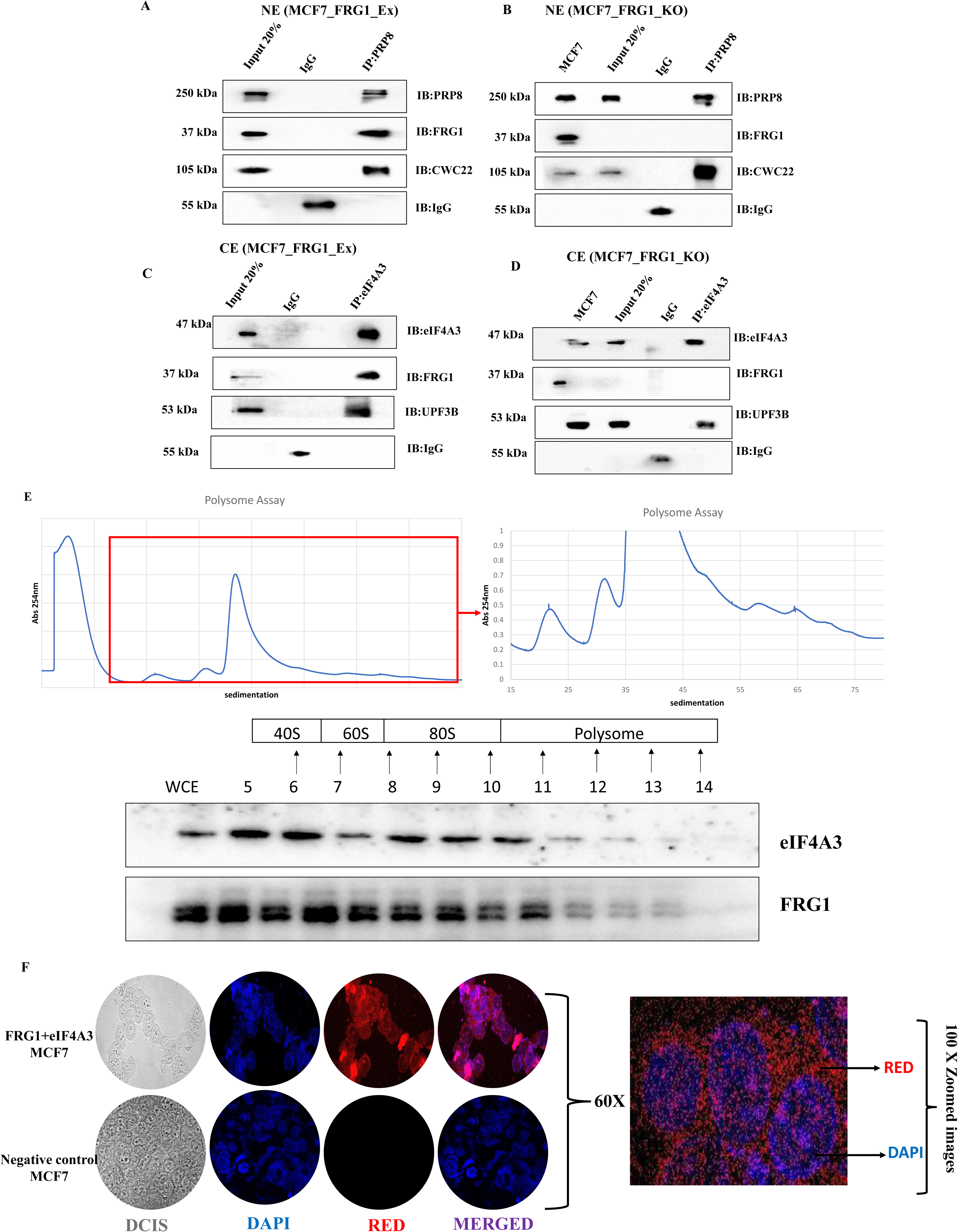
Interaction of FRG1 with Spliceosome and Exon Junction Complex. **(A-B)** Representative Western blots demonstrating co-immunoprecipitation with the PRP8 antibody in the nuclear extracts (NE) of MCF7 cells. Panel **(A)** shows results from cells with FRG1 overexpression, while panel **(B)** displays findings from cells with FRG1 knockdown. Co-immunoprecipitation with IgG was used as a negative control. **(C-D)** Representative Western blots illustrating co-immunoprecipitation with the eIF4A3 antibody in the cytoplasmic extract (CE) of MCF7 cells with ectopic FRG1 expression **(C)** and reduced FRG1 expression **(D)**. Co-immunoprecipitation with IgG was used as a negative control. **(E)** Upper panel: Sucrose gradient fractionation of cytoplasmic extracts from MCF7 cells. The polysome profile displays relative absorbance at 254 nm across a 20–40% sucrose gradient, Lower panel: Immunoblot analysis of gradient fractions using antibodies against FRG1 and eIF4A3. Immunoblot lanes corresponding to the 40S, 60S, 80S ribosomal subunits, and polysome peaks are indicated. **(F)** Proximity Ligation Assay (PLA) performed using antibodies against FRG1 and eIF4A3. Representative confocal images of MCF7 cells (acquired at 60× magnification) are shown in the upper panel. Red puncta indicate positive PLA signals, representing protein–protein proximity (<40 nm), suggestive of a potential interaction. Nuclei are counterstained with DAPI (blue). The lower panel shows 100× magnified views of the PLA puncta, emphasizing the specificity of the observed interaction.

Next, we examined whether FRG1 affects the architecture of the EJC. We focused on eIF4A3, a core EJC component [72], and UPF3B, a key factor in the NMD pathway. Co-IP using an anti-eIF4A3 antibody on cytoplasmic extracts of FRG1-perturbed cells revealed interactions between eIF4A3, FRG1, and UPF3B (Fig. 7 C). Notably, in FRG1-deficient cells, the interaction between eIF4A3 and UPF3B was maintained (Fig. 7 D), indicating that FRG1 is not essential for EJC complex stability. To further probe the potential role of FRG1 in post-transcriptional regulation, we performed polysome sucrose gradient fractionation followed by immunoblotting. Remarkably, FRG1 co-sedimented with eIF4A3 across polysome fractions (Fig. 7 E), suggesting a previously unrecognized association of FRG1 with actively translating ribosomal complexes. This observation strongly supports a functional link between FRG1 and the translation-coupled phase of EJC activity, hinting at a novel regulatory role for FRG1 in mRNA surveillance or translation.

To validate the physical proximity suggested by co-sedimentation, we conducted a Proximity Ligation Assay (PLA) to test whether FRG1 and eIF4A3 are spatially close enough to imply direct or complex-mediated interaction. The PLA revealed distinct and abundant red fluorescent puncta in cells, confirming that FRG1 and eIF4A3 are localized within less than 40 nm of each other—a range consistent with direct protein-protein interaction (Fig. 7 F). These compelling findings strongly support the conclusion that FRG1 either directly binds eIF4A3 or is stably associated within the same molecular complex. Altogether, this series of experiments unveils an exciting new aspect of FRG1 biology and firmly establishes its integration into key post-transcriptional gene regulatory pathways.

## Discussion

Research involving FRG1 has predominantly focussed on its association with FSHD and muscle function [40,42,73]. Over the past decade, there has been a substantial increase in understanding the additional roles of FRG1 in tumorigenesis and angiogenesis [43–46]. Although few studies have indicated that FRG1 expression levels affect the RNA biogenesis of specific genes, but the exact mechanism is unknown [40,47,74]. The results presented in this study sheds light on FRG1’s role in modulating NMD processes, and its underlying molecular mechanism suggesting the need for maintaining its optimal levels.

Our previous finding revealed that FRG1 depletion decreases the expression levels of NMD factors such as MAGOH, PININ, ROD1, UPF3B, SMG1, and ICE1 [52]. Another study demonstrated that ICE1 depletion reduces NMD efficiency, indirectly suggesting a potential role for FRG1 in regulating NMD [38]. We observed that reduced FRG1 increased expression level of UPF1 transcriptionally as well as translationally, thereby increasing NMD efficiency. Thus, it is possible that FRG1, acting as a dual regulator of NMD genes may play a role in stabilizing the NMD pathway, ensuring its robustness in quality control and regulating normal gene expression. Previously, multiple studies have highlighted the role of NMD factors in maintaining cellular homeostasis. The first indication of a buffering mechanism came from a microarray profiling analysis on NMD deficient HeLa cells with depleted levels of the NMD factor UPF1. In this, amongst the 195 significantly upregulated transcripts, mRNA encoding the NMD factor SMG5 was identified [1]. In yet another study, SMG5 mRNA was found to be regulated by the UPF3B-independent pathway [32]. Huang et al. demonstrated the presence of a conserved feedback regulatory network wherein depletion of UPF1 in HeLa cells resulted in upregulation of NMD factors-UPF2, UPF3B, SMG1, SMG5, SMG6, and SMG7 [75]. Study by Yepiskoposyan et al. showed that long 3’UTRs of NMD substrate mRNAs (UPF1, SMG6, and SMG7) trigger NMD, demonstrating autoregulation of NMD in human cells [76]. From all of these studies, we may speculate that along with the aforementioned NMD factors, FRG1 might be a part of the feedback regulatory network, preserving the integrity of NMD. NMD feedback loops are far more intricately regulated than currently understood. In future, genome-wide analysis may help elucidate new NMD factors that might provide versatile and subtle feedback control to achieve optimal NMD activity and protect NMD from perturbation.

DUX4 is a well-established transcriptional regulator of FRG1 [64,66]. Previously Feng et al. reported that DUX4 reduces NMD efficiency triggering proteolytic degradation of UPF1 [65]. Consistent with this, our observation revealed that NMD efficiency is directly, but inversely modulated by FRG1. We ascertained that DUX4-mediated NMD inhibition is regulated by FRG1 expression. Delving into its molecular intricacy, we have shown that FRG1 mediates degradation of UPF1, which results in NMD inhibition. Thus, the close coupling between DUX4 and FRG1 expression levels might be providing a regulatory mechanism to maintain NMD homeostasis. However, it is yet to be determined if increased UPF1 protein levels are primarily responsible for FRG1-induced NMD regulation or merely one of the several contributing factors. Nonetheless, the expression levels of FRG1 in altering NMD efficiency surpassing DUX4 may prove useful to gain insight into the biological relevance of this mechanism. It is also tempting to speculate that FRG1-induced dysregulation of the ubiquitin-proteasome system is responsible for triggering UPF1 protein degradation. The precise mechanism by which FRG1 induces UPF1 proteolysis or whether FRG1 itself acts as E3 ubiquitin ligase remains to be elucidated.

FRG1 serves as a structural component of spliceosome C complex by interacting with PRP8 [51] and previous studies have linked increased FRG1 expression with abnormal splicing [48,77]. Herein, we aimed to investigate whether it regulates NMD by being a structural component of the Spliceosomal C complex. In this study, using FRG1 perturbed cell lines, carrying out co-immunoprecipitation assays, we have established it as a structural component of the Exon Junction Complex and Spliceosome Complex. However, we could also find that FRG1’s absence did not disrupt the interaction between CWC22 (a spliceosome molecule) and PRP8 or eIF4A3 (core EJC molecule) and UPF3B. However, in this context, the finer alterations in the spliceosome or exon junction complexes remain unexplored, leaving open the possibility that FRG1 may influence their functionality. Alternatively, other NMD factors might be compensating for the loss of FRG1 to maintain the structural integrity of the complexes. This requires further validation. A comprehensive mass spectrometry analysis conducted by Singh et al. [78] using immunoprecipitation from HEK293 cells expressing FLAG-tagged eIF4A3 and mFLAG-Magoh, revealed a broad enrichment of EJC core and peripheral components, along with spliceosomal proteins, heterogeneous nuclear ribonucleoproteins (hnRNPs), ribosomal subunits, and serine/arginine-rich (SR) proteins. These findings underscore the complex and multifunctional nature of the EJC and its dynamic associations within the mRNP environment. In line with this, our polysome profiling data provide compelling evidence for a potential interaction between FRG1 and eIF4A3, strongly suggesting that FRG1 may associate with the EJC. This association further supports the emerging view of FRG1 as a participant in post-transcriptional gene regulatory pathways, possibly contributing to the coupling of splicing, translation, and mRNA surveillance mechanisms. Future studies will be needed to address the degree of FRG1-dependent interplay between splicing and NMD pathways.

In summary, this study provides mechanistic evidence that FRG1 inversely regulates the NMD machinery. Crucially, we demonstrated that FRG1 directly modulates the NMD pathway by influencing the ubiquitination of UPF1, an effect that surpasses the regulatory impact of DUX4. The identification of FRG1 as an upstream regulator of NMD genes offers valuable insights into the mechanisms underlying the fine-tuning of NMD. The findings from this study pave the way for the development of novel therapeutic strategies targeting diseases associated with FRG1 dysregulation.

## Abbreviations

FRG1: FSHD Region Gene 1
UPF1: Up-frameshift Protein 1
DUX4: Double homeobox 4
PRP8: Pre-mRNA processing factor 8
CWC22: Spliceosome associated protein homolog
UPF3B: Up-frameshift Protein 3B
ICE1: Interactor of little elongation complex ELL subunit 1
MAGOH: Mago Homolog
EJC: Exon junction complex
SURF: SMG-1-Upf1-eRF1-eRF3 complex
GAPDH: Glyceraldehyde 3-phosphate dehydrogenase
DECID: Decay-inducing complex
SMG: PI3K related kinase
ROD1/PTBP3: Polypyrimidine Tract Binding Protein 3
Rbfox1: RNA-binding fox-1 homolog 1
FXR1: FMR1 autosomal homolog 1

## Acknowledgement

Not Applicable

## Author Contribution

A.P.: performed Western Blot, qRT-PCR, luciferase reporter assay, cell culture, co-immunoprecipitation, zebrafish experiments, formal analysis of data, writing-original draft and editing. T.M. performed cell culture, western blotting, co-immunoprecipitation, MG132 inhibitor treatments, and polysome assays, proximity ligation assay and contributed to editing the manuscript draft. A.S. and S.T.: analyzed RNA seq data. S.S.M.: helped in zebrafish genotyping, maintaining zebrafish lines, and imaging work. R.K.S.: designed zebrafish-related experiments, carried out microinjection in zebrafish embryos, and reviewed the manuscript. M.D.: conceptualized the study, analysed the data, wrote and reviewed the manuscript, and acquired the funding.

## Ethics Approval

All zebrafish-related experiments were performed following the protocol approved by Institutional Animal Ethics Committee (IAEC), ILS, Bhubaneswar under the protocol number ILS/IAEC-213-AH/APR-21 and ILS/IAEC-364-AH/SEPT-24.

## Data Availability Statement

Data associated with this work are available upon request from the corresponding author.

---

**Supplementary Figure 1. Screening of frg1 knockout AB strain zebrafish. (A-B)** Agarose gel images showing amplified frg1 exon-1(153 bp) in wild type (WT), heterozygous (HETERO), and homozygous (HOMO) zebrafish embryos **(A)** Agarose gel image showing amplified frg1 exon-1(153 bp) in wild type (WT) but not in homozygous (HOMO) zebrafish embryos. **(C-F)** show image of zebrafish embryos having expression of frg1 in wild type (WT) **(C)** and frg1 knockout (frg1) **(D)** conditions at 24 hours post fertilization (hpf) stage. **(E-F)** show image of zebrafish embryos having expression of frg1 in wild type (WT) **(E)** and frg1 knockout (frg1) **(F)** conditions at 48 hours post fertilization (hpf) stage.

**Supplementary Figure 2. NMD-sensitive isoform levels upon FRG1 knockdown.** The figure shows the isoform expression profile plots displaying isoform structures in the upper panel along with the annotated ensembl IDs. The lower panel displays isoform expression and isoform fraction of NUB1 **(A)**, ACOX1 **(B)**, XPO5 **(C)**, ZMYND11 **(D)**, LZTFL1 **(E)**, SLMAP **(F)**, RAD51D **(G)** and TRIM47 **(H)** undergoing isoform switching upon FRG1 depletion (FRG1_KD vs control). Red box indicates the NMD-sensitive isoforms of the corresponding genes.

**Supplementary Figure 3. NMD-sensitive isoform levels upon FRG1 knockout.** The figure shows the isoform expression profile plot displaying isoform structures in the upper panel along with the annotated ensembl IDs. The lower panel displays isoform expression and isoform fraction of ERGICI1 **(A)**, ZW10 **(B)**, OSBPL7 **(C)**, REEP4 **(D)**, MRPL22 **(E)**, DTWD2 **(F)**, OAS2 **(G)**, DRAM1 **(H)**, BMERB1 **(I)** and WDR20 **(J)** undergoing isoform switching in the absence of FRG1 (FRG1_KO vs control). Red box indicates the NMD-sensitive isoforms of the corresponding genes.

