## Supplementary figures and images for "FRG1 Regulates Nonsense-Mediated mRNA Decay by Modulating UPF1 Levels"

### Supplementary Figure 1

##### Supplementary Figure 1

A

F1 generation

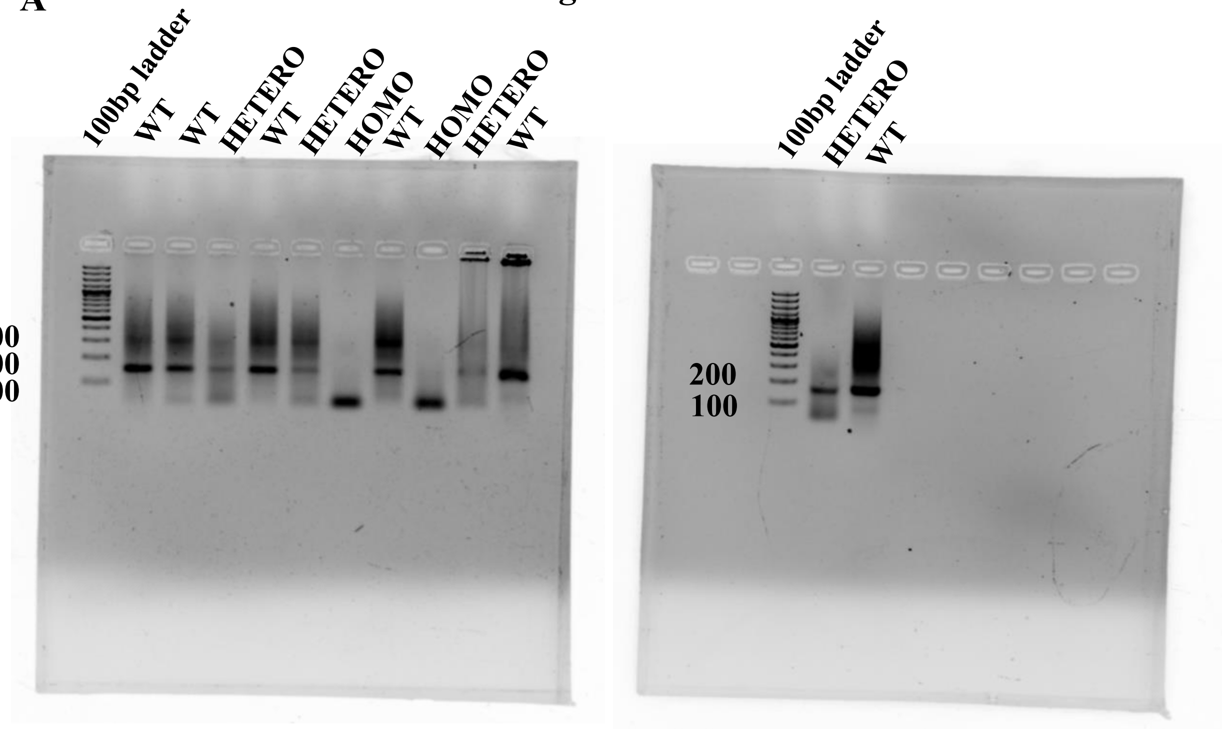


B

F2 generation

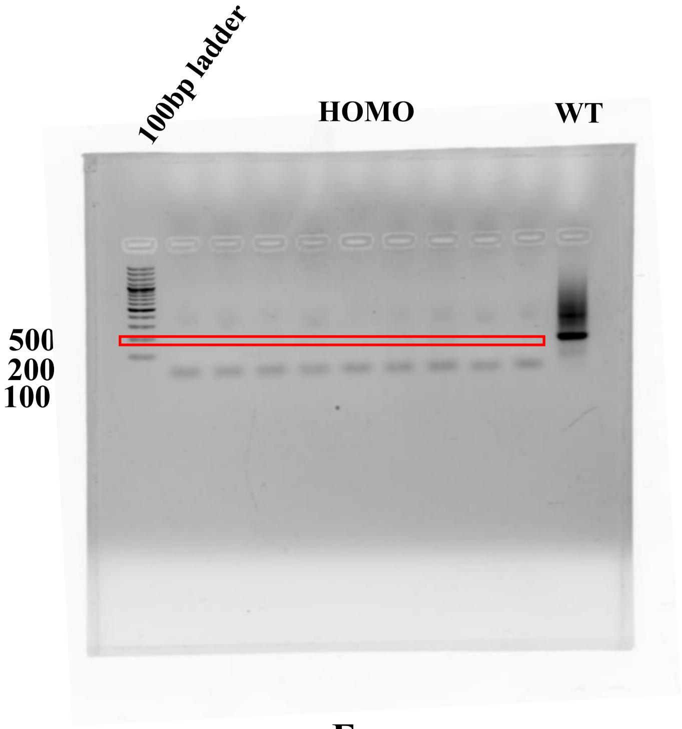


C

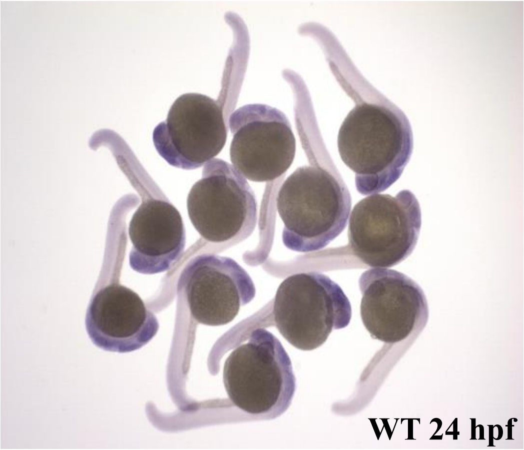


D

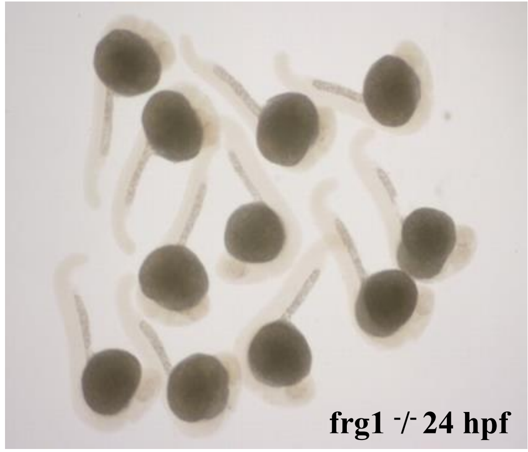


E

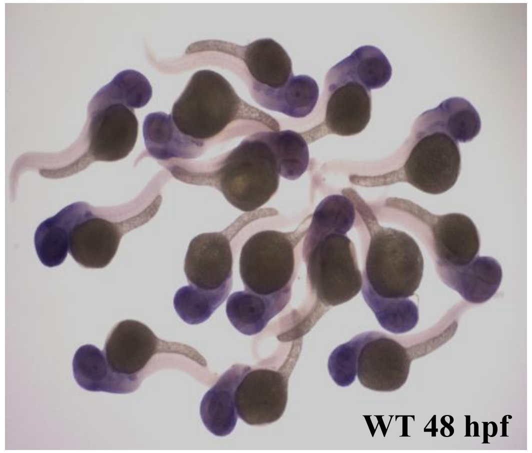


F

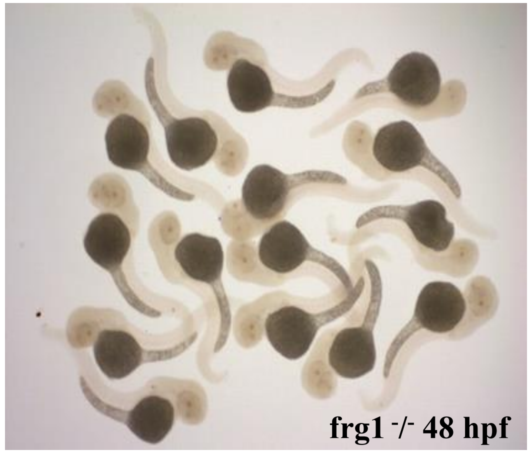

### Supplementary Figure 2

#### Supplementary Figure 2

A

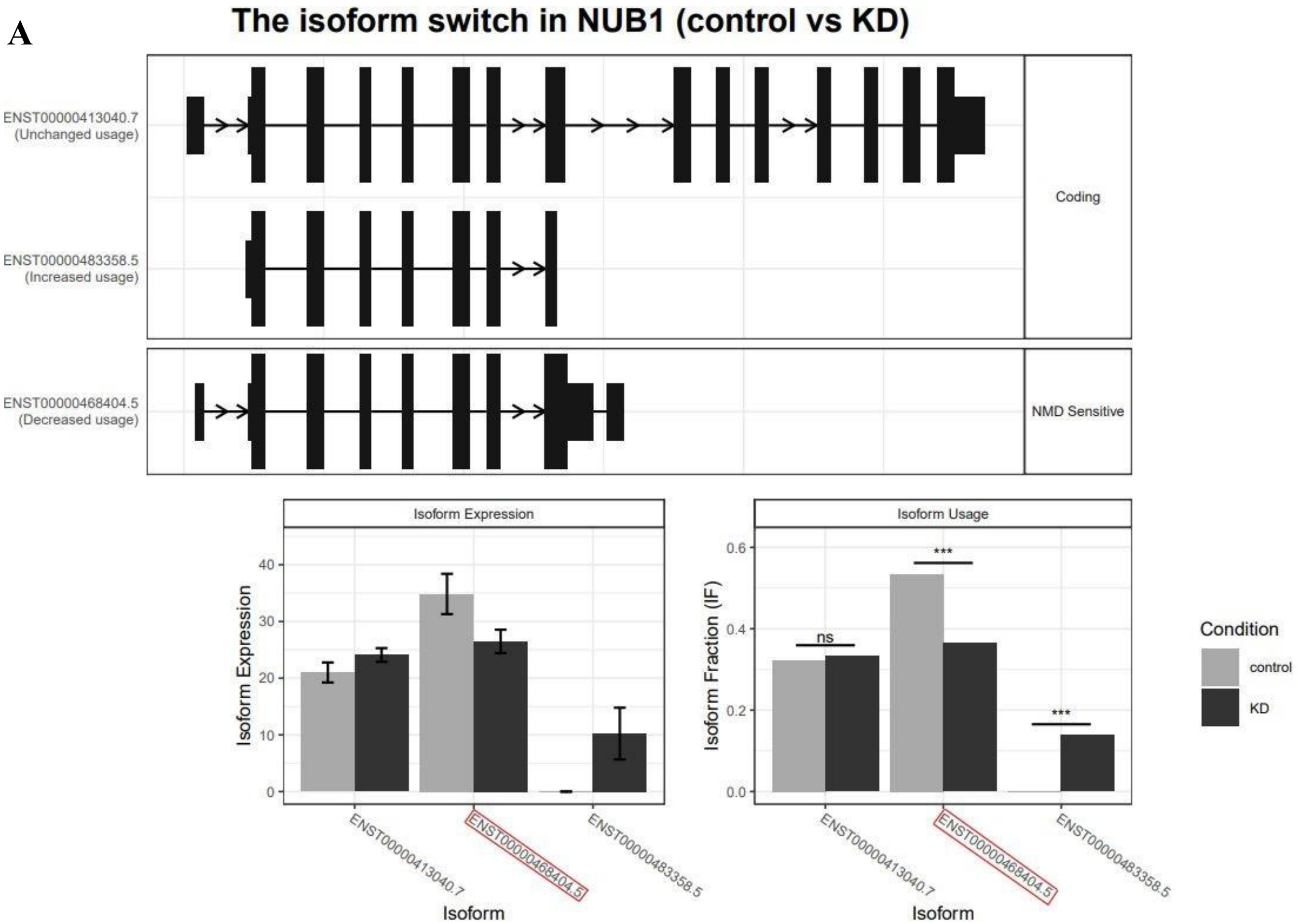

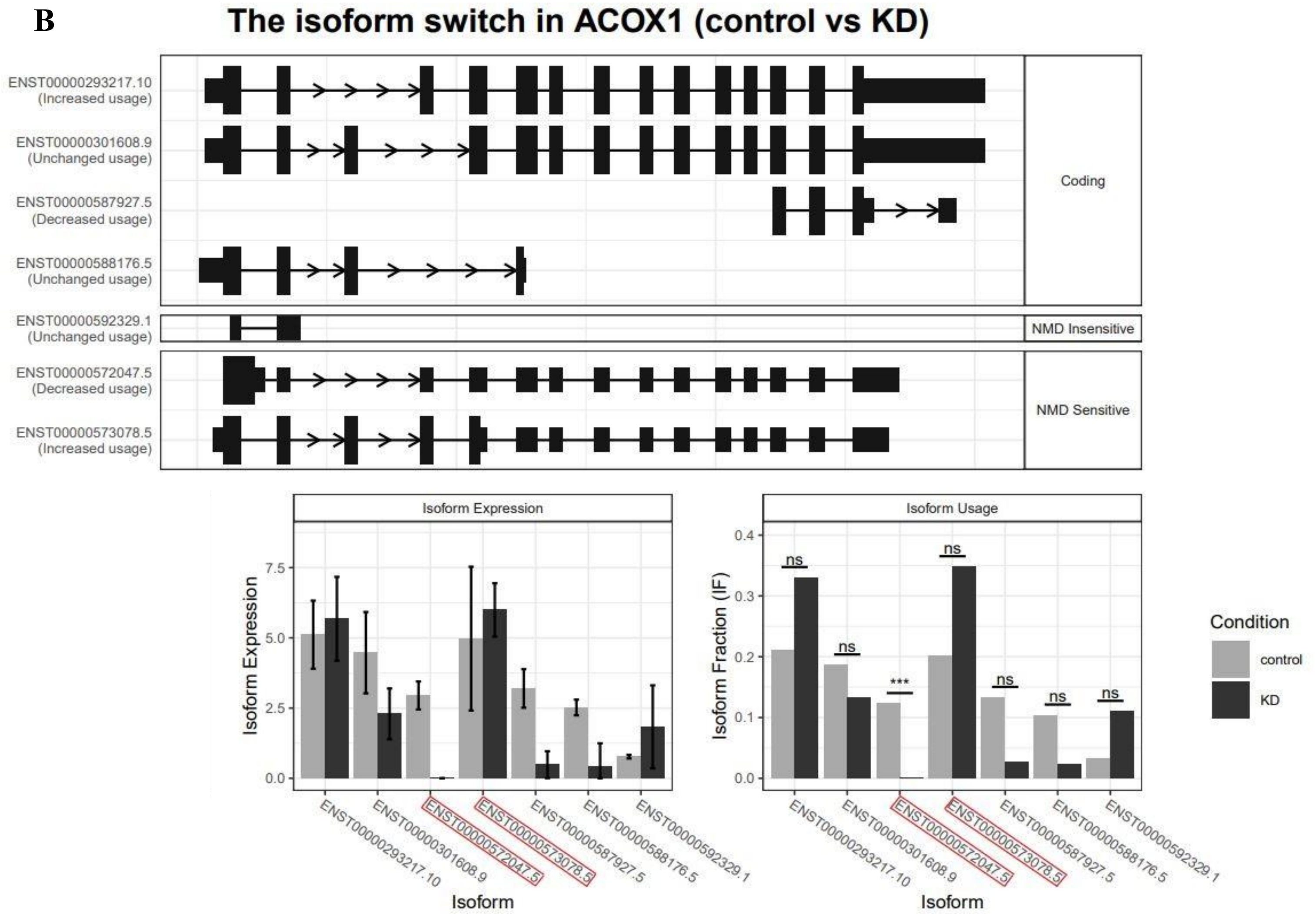

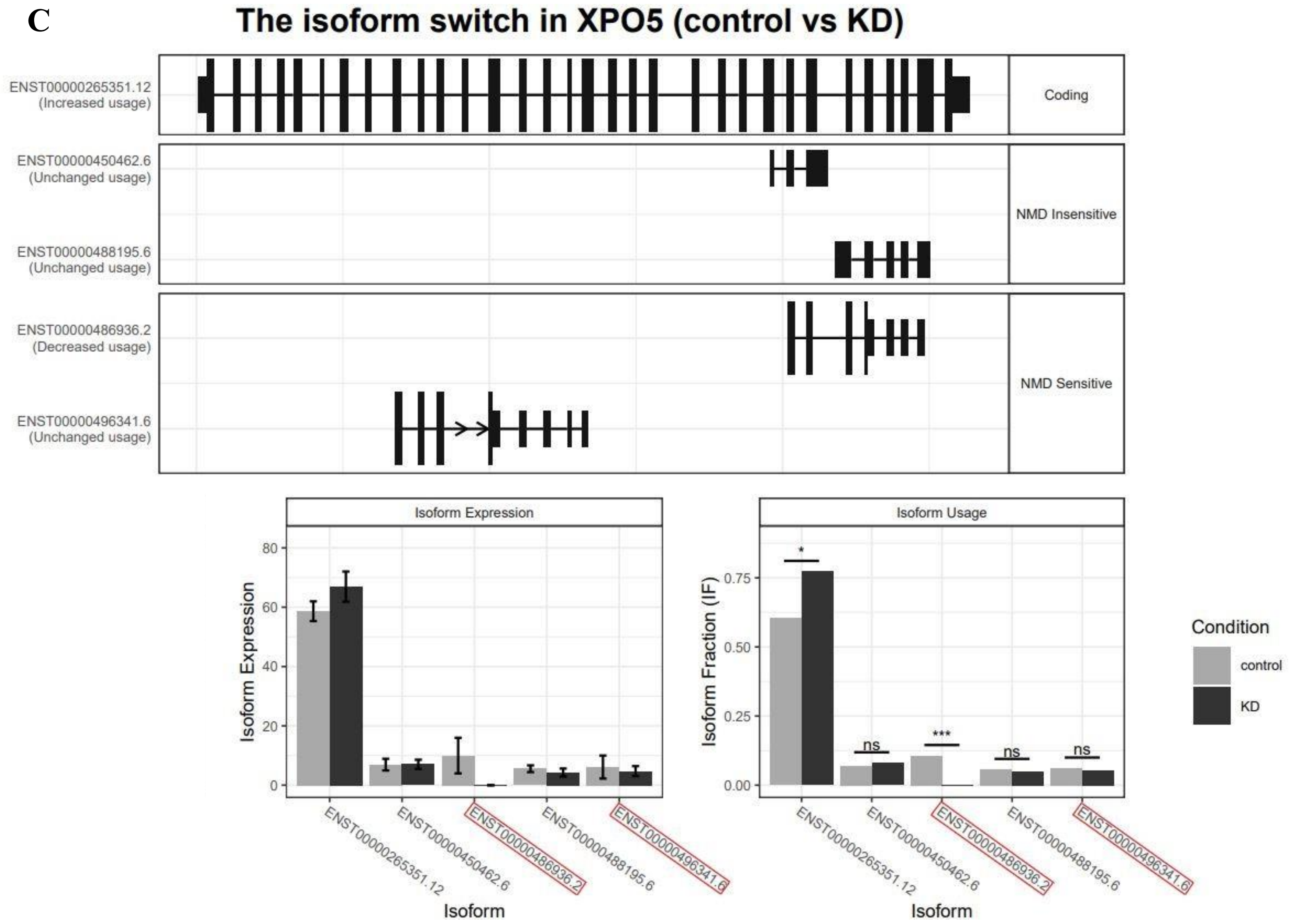

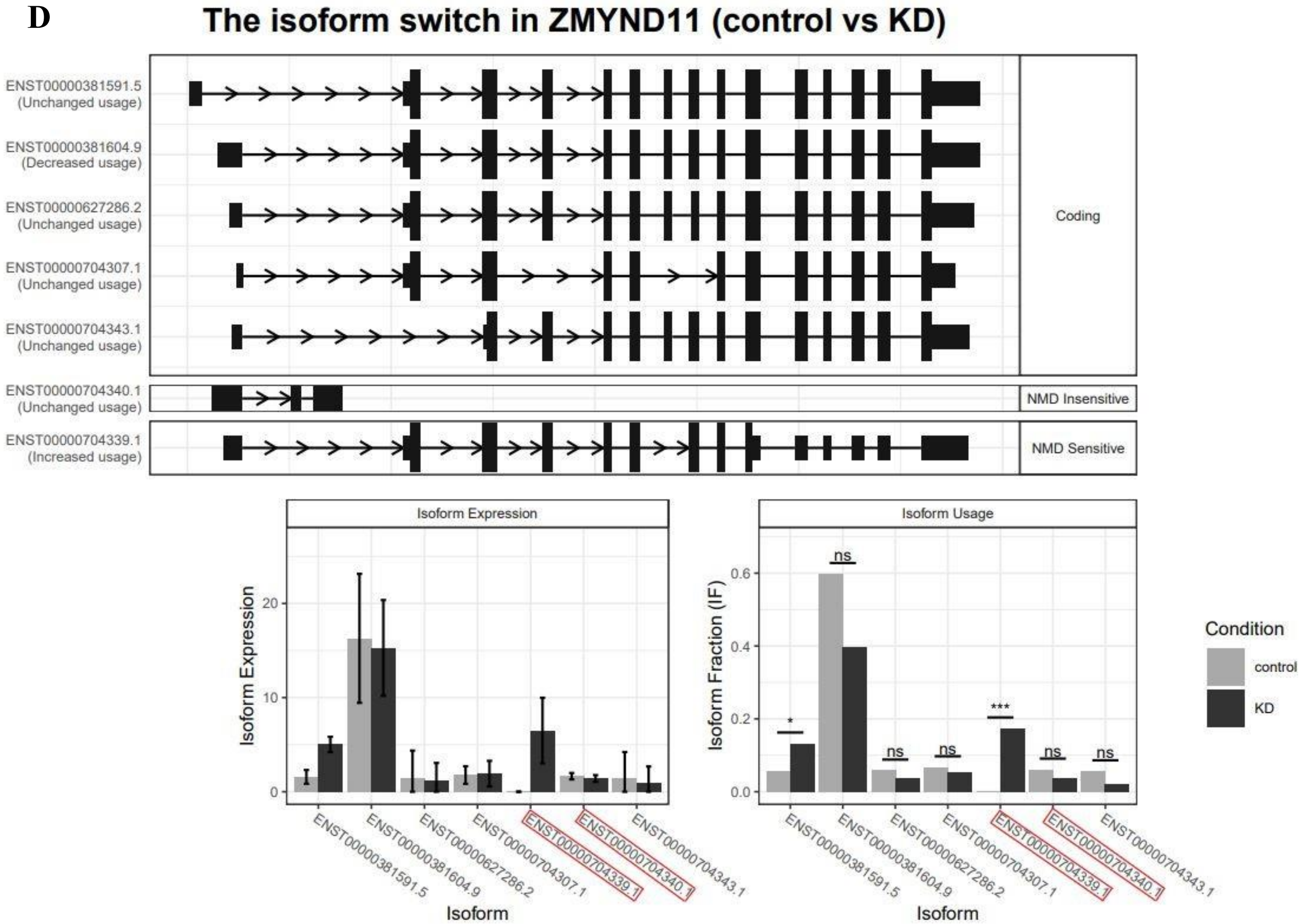

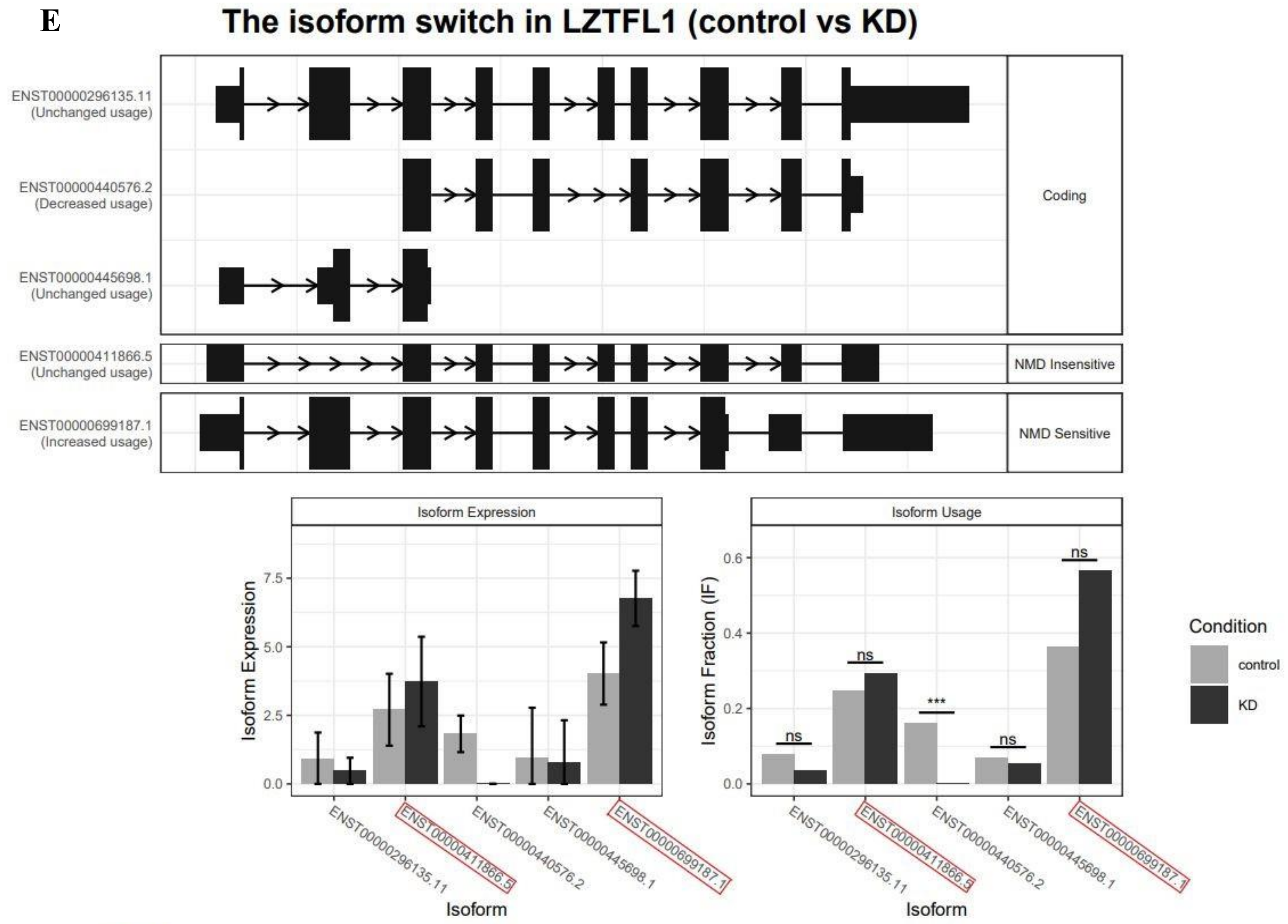

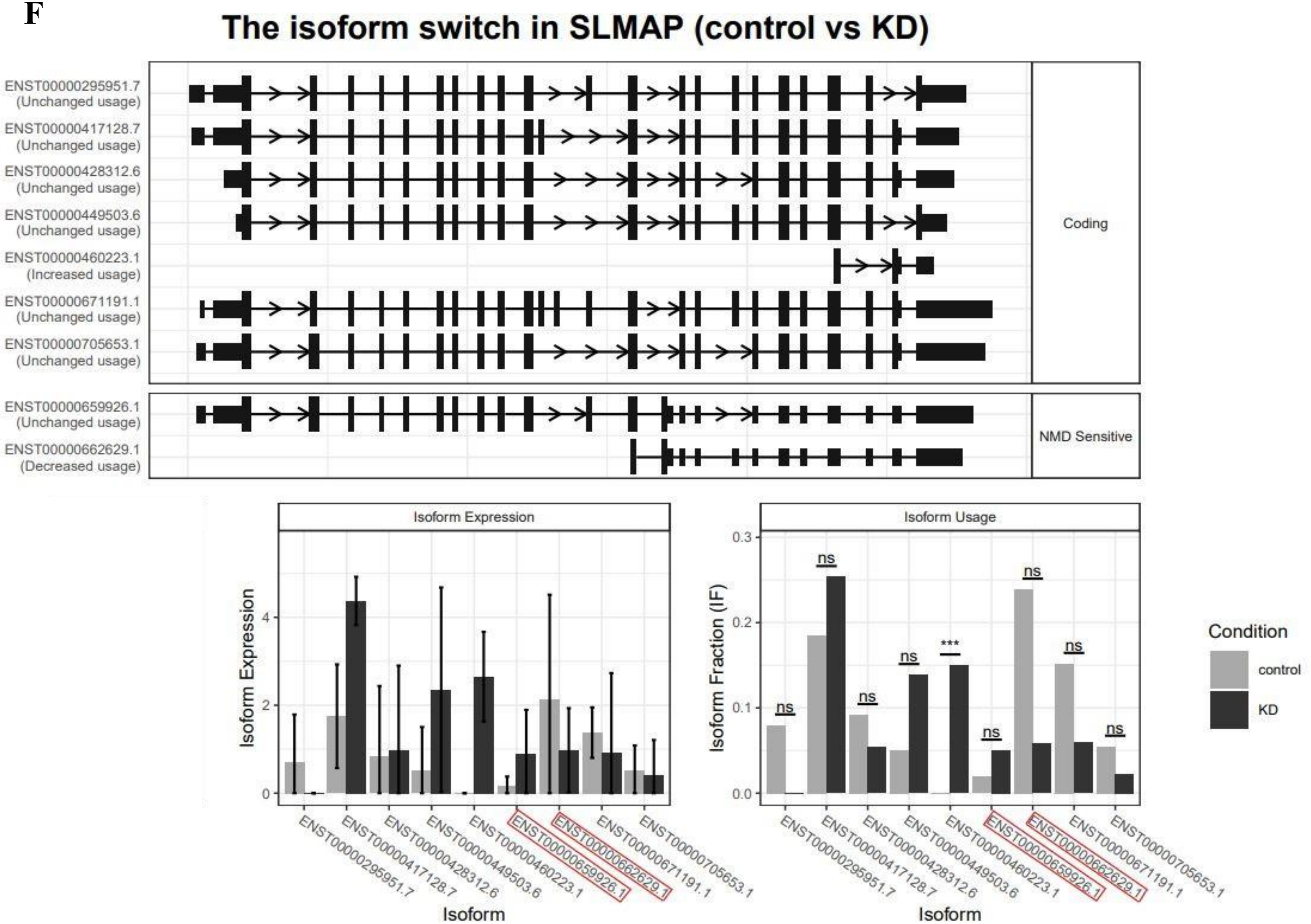

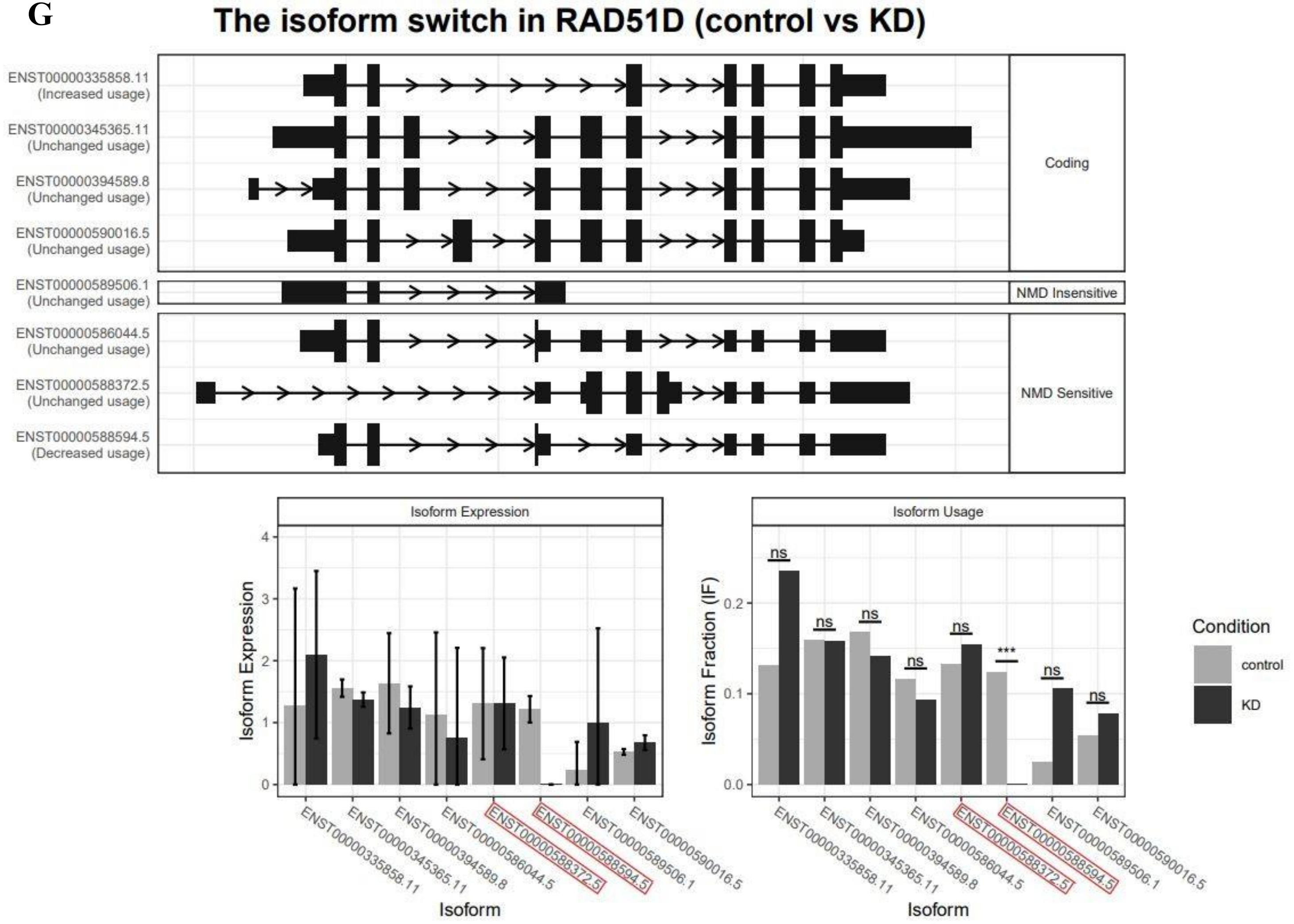

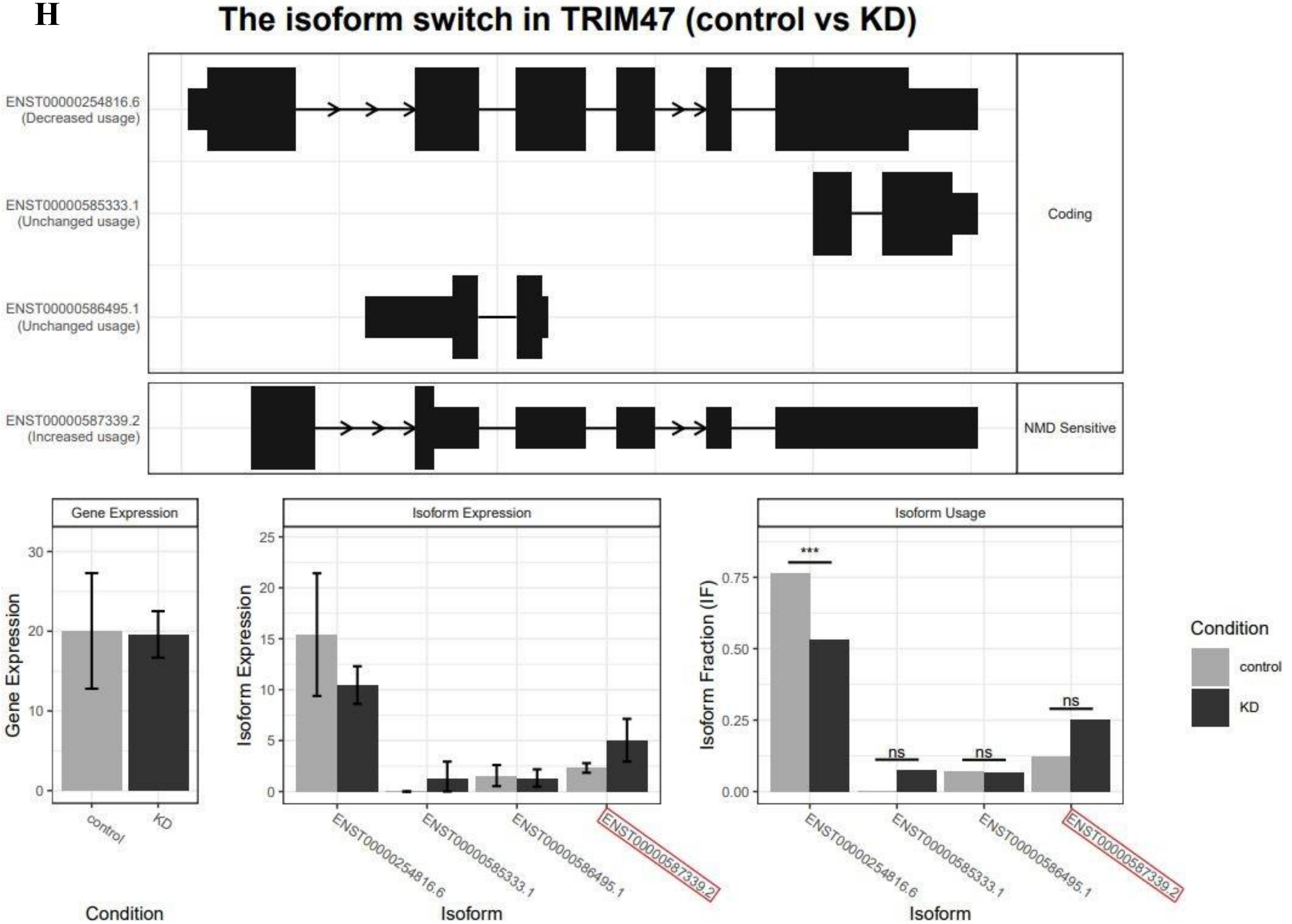

### Supplementary Figure 3

**Supplementary Figure 3**

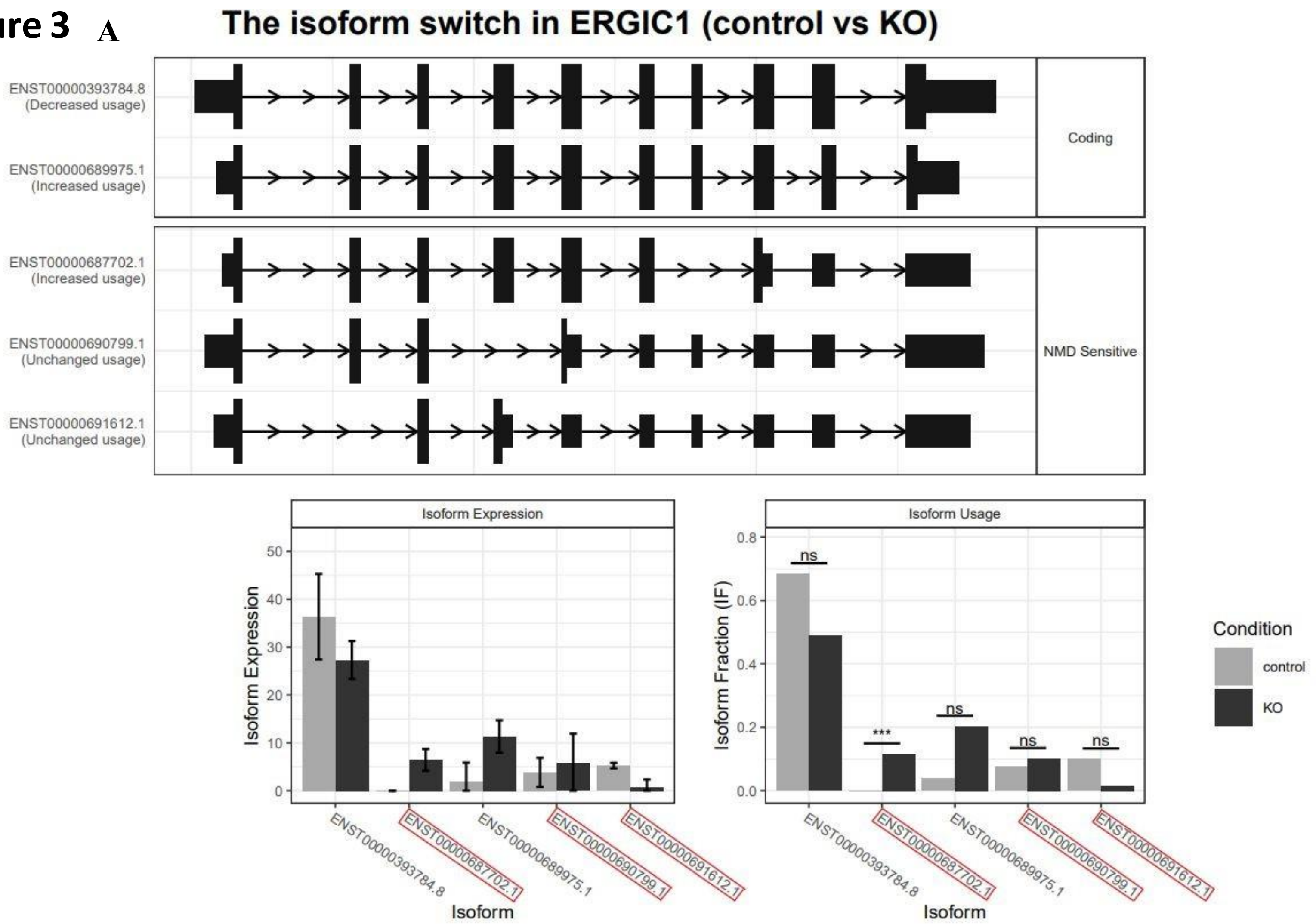

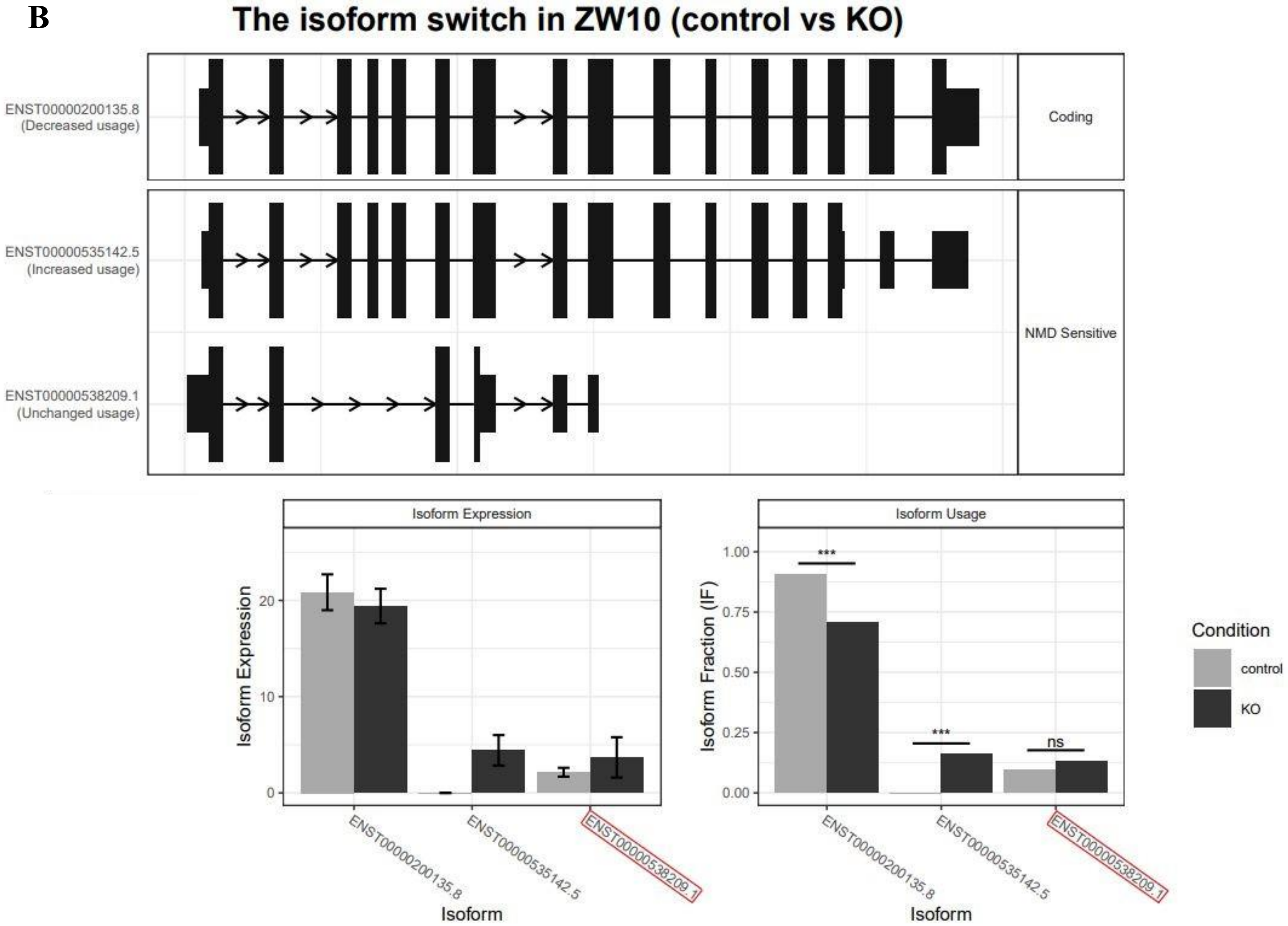

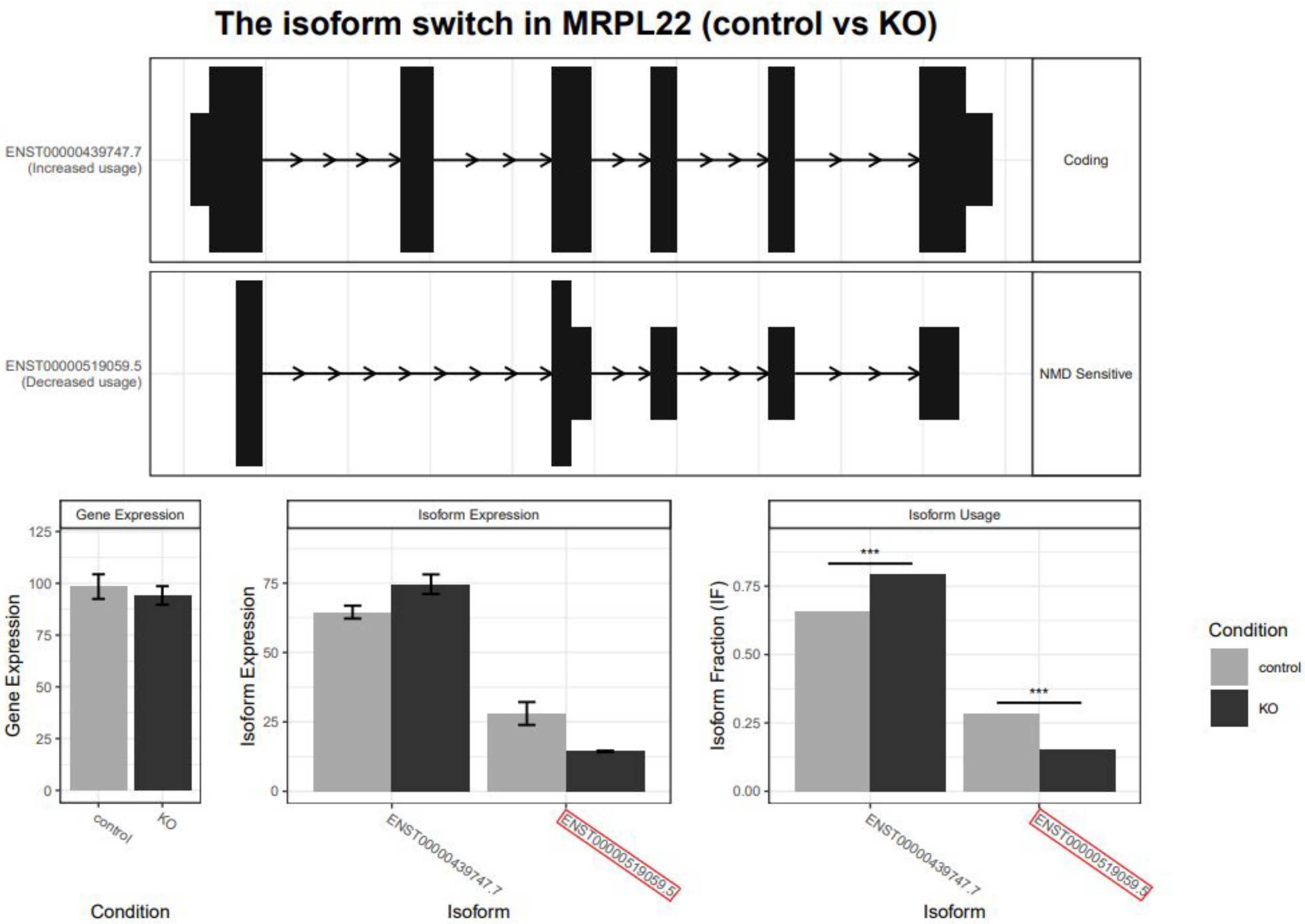

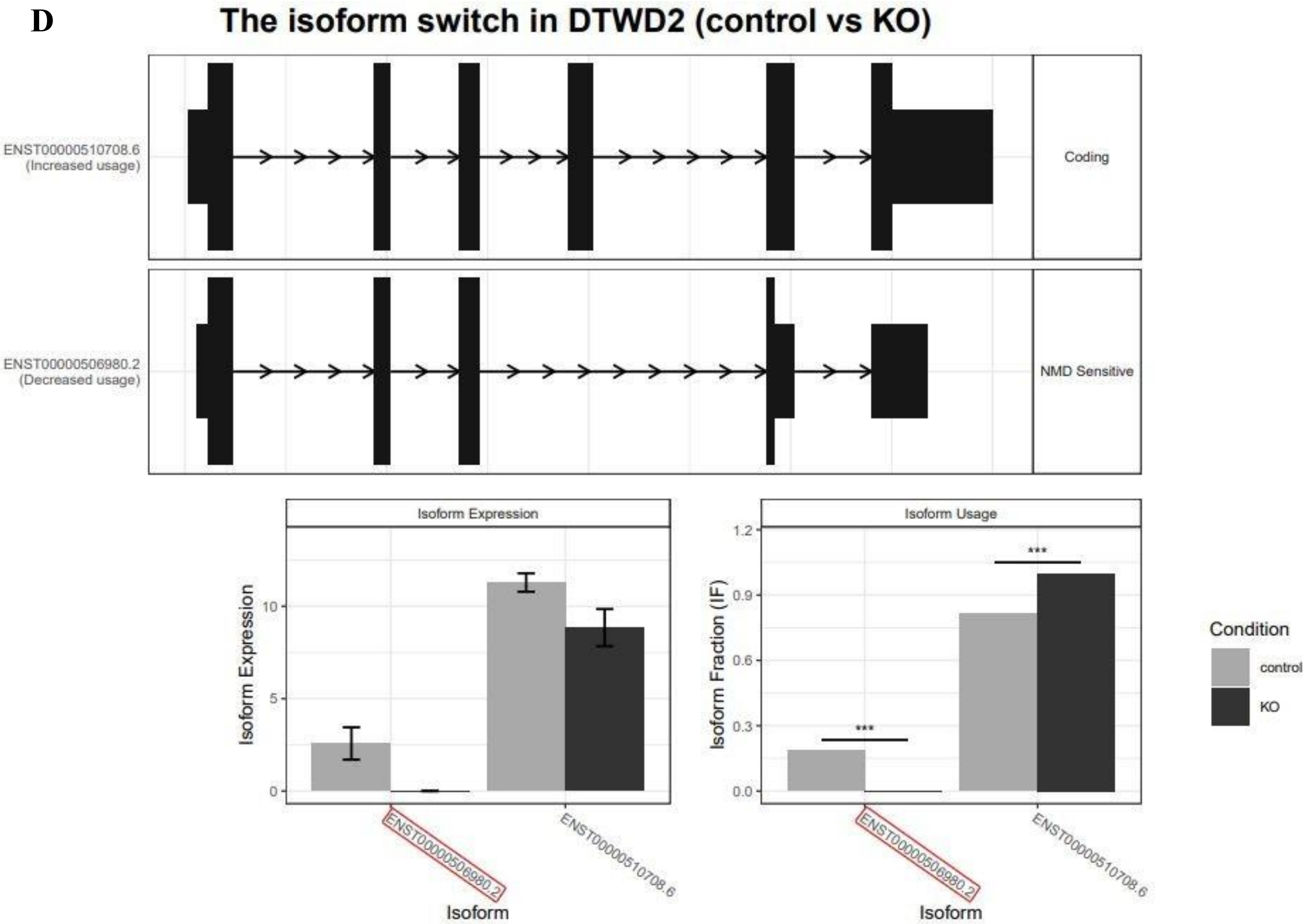

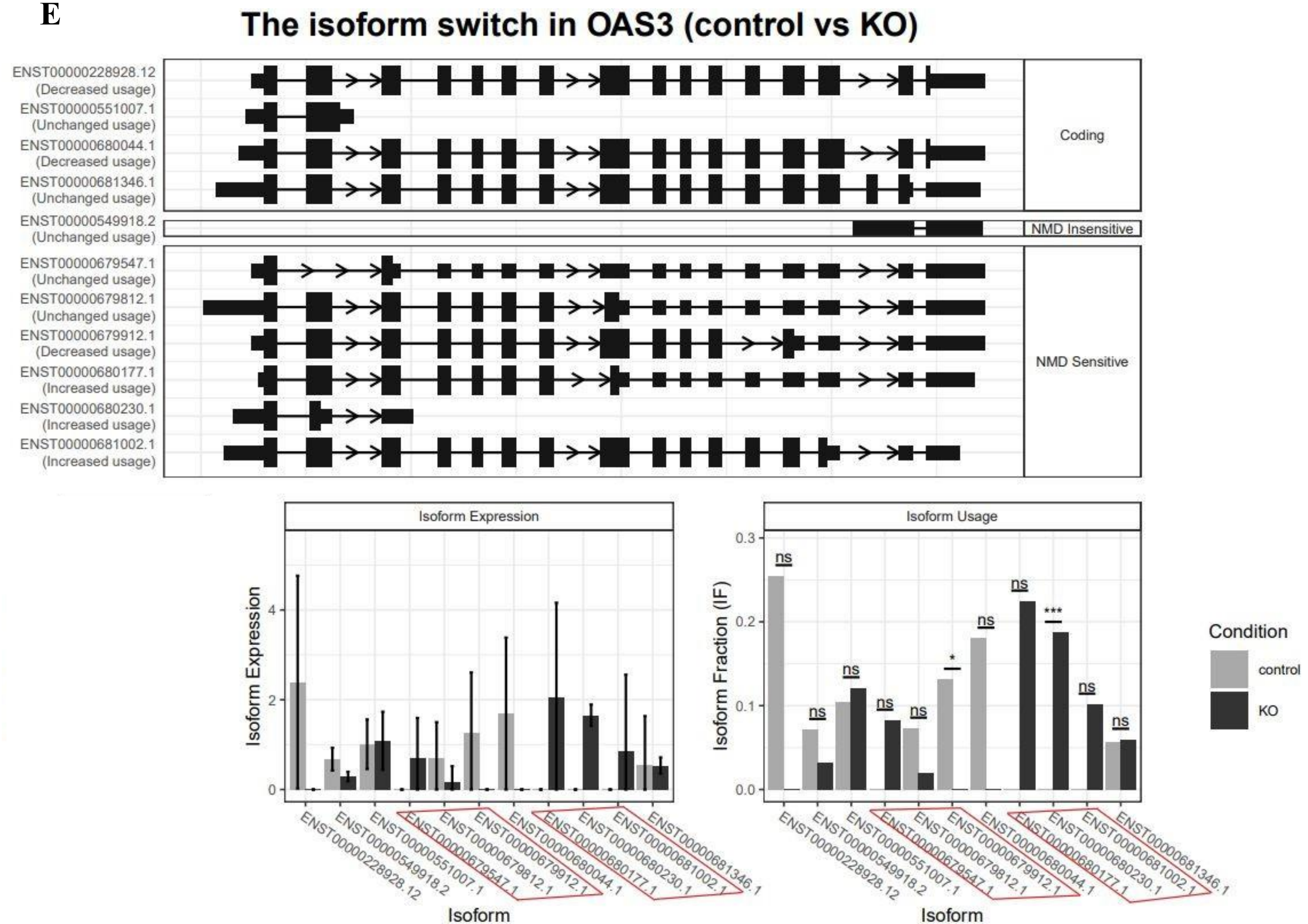

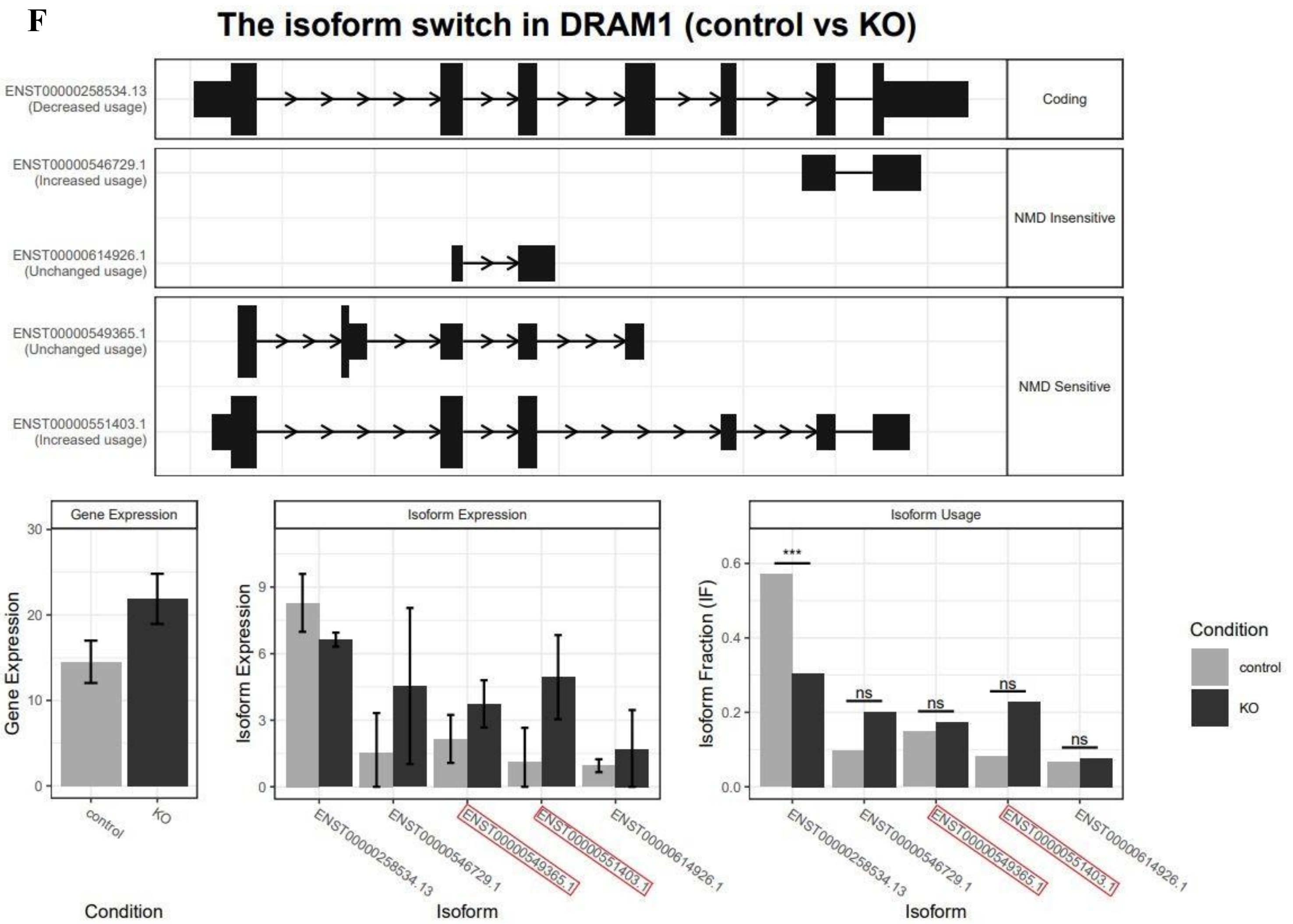

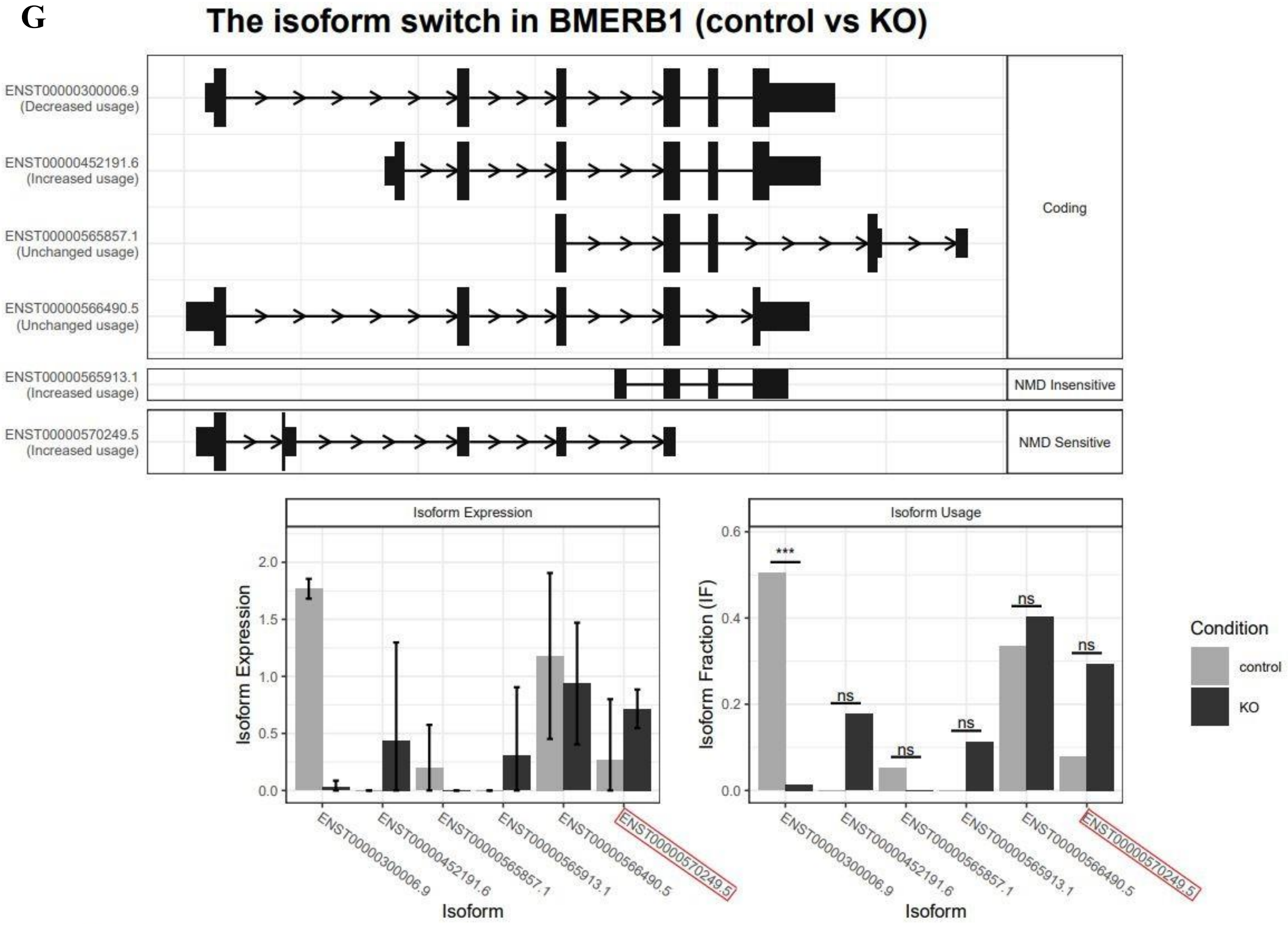

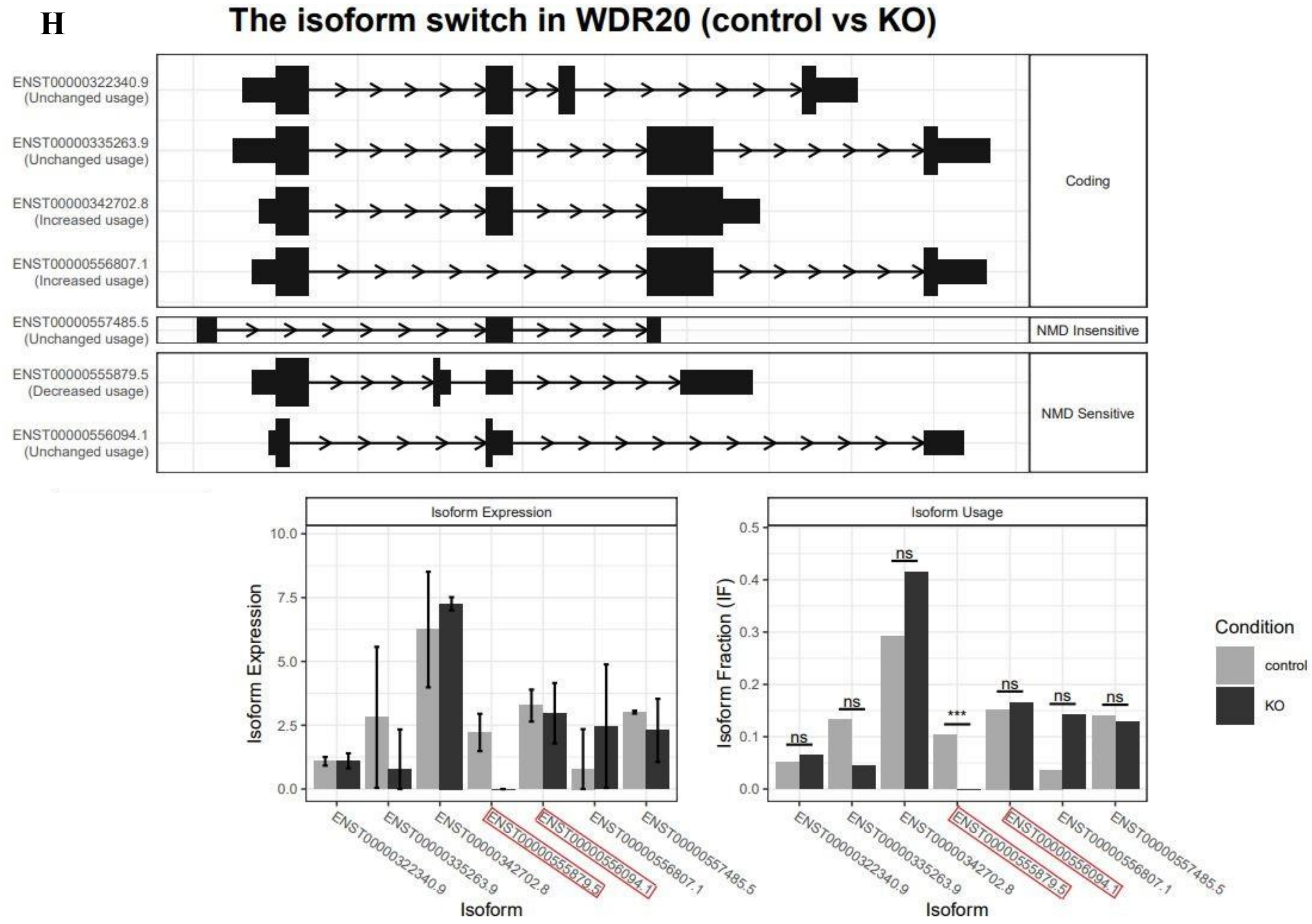
