## Supplementary Tables for "FRG1 Regulates Nonsense-Mediated mRNA Decay by Modulating UPF1 Levels"

### Table S1: List of oligos used in this study

| PURPOSE | SEQUENCE 5'to3' |
|---|---|
| **UPF1 qRT-PCR** | F: ATATGCCTGCGGTACAAAGG<br>R: AGCTCAATGGCGATCTCATC |
| **GAPDH qRT-PCR** | F: ACCCAGAAGACTGTGGATGG<br>R: TCTAGACGGCAGGTCAGGTC |
| **ZF UPF1 qRT-PCR** | F: GTGGTTCGTCTGTGTGCTAA<br>R: CATGTTGCGGATCTGATTGTG |
| **ZF EF1A qRT-PCR** | F: ATCACCAAGGAAGTCAGCG<br>R: ATCTTCCATCCCTTGAACCAG |
| **ZF WISH frg1 PRIMERS** | F: ATTGTTGGAGGGTGGTGGAC<br>R: AGCAGTGCCTCATGGAACTT |
| **ZF frg1 exon-1** | F: AGTGTTGGGTCTGTGTGTGATC<br>R: GAAACACAGCTCAACGTCAAAC |
| **ZF frg1 3'UTR** | F: CTTTGGAGAAGCCTGTGTCACT<br>R: ATTCTGGAAGCTAGTCTCTATC |
| **ZF frg1_sgRNA (EXON-1)** | GAACCTTTCAGGACCAGTTTGG |
| **ZF frg1_sgRNA (EXON- 9)** | GGCCTGACTGTATATCTGCAGGG |

### Table S2: List of sources

| Reagent/Resource | Source | Identifier |
|---|---|---|
| Zebrafish: *Danio rerio* (AB strain) | ZIRC / Laboratory stock | RRID:ZFIN_ZDB-GENO-960809-7 |
| E3 embryo medium reagents (NaCl, KCl, $CaCl_2 \cdot 2H_2O$, $MgCl_2 \cdot 6H_2O$) | Sigma-Aldrich | Cat# S9888 (NaCl), P5405 (KCl), C7902 ($CaCl_2 \cdot 2H_2O$), M2670 ($MgCl_2 \cdot 6H_2O$) |
| Sodium hypochlorite | Sigma-Aldrich | Cat# 425044 |
| Recirculating water system | Techniplast, India | N/A |
| sgRNA design method | Varshney et al., 2015 | DOI:10.1038/nprot.2015.050 |
| gRNA scaffold oligo | Integrated DNA Technologies (IDT) | Custom synthesis |
| MEGAscript™ SP6 Transcription Kit | Invitrogen | Cat# AM1330 |
| MEGAscript™ T7 Transcription Kit | Invitrogen | Cat# AM1334 |
| mMESSAGE mMACHINE™ Kit | Invitrogen | Cat# AM1344 |
| pCS2-nCas9n nanos 3′UTR plasmid | Addgene | Plasmid #62542 |
| XbaI-HF restriction enzyme | New England Biolabs (NEB) | Cat# R0145S |
| Phusion High-Fidelity DNA Polymerase | Thermo Scientific | Cat# F530S |
| Femtojet microinjector | Eppendorf | Cat# 5247 000.013 |

### Table S3: List of software sources

| Software/Algorithm | Source | Identifier |
|---|---|---|
| ImageJ (FIJI) | NIH | RRID:SCR_003070 |
| GraphPad Prism (v9 or v10) | GraphPad Software | RRID:SCR_002798 |
| R statistical software (v4.x) | R Foundation | RRID:SCR_001905 |
| CRISPR design tool | CHOPCHOP | RRID:SCR_015723 |

### Table S4: List of antibodies sources

| **Reagent/Resource** | **Source** | **Identifier** |
|---|---|---|
| Rabbit recombinant monoclonal anti-FRG1 (clone EPR13098) | Abcam (MA, USA) | Cat# ab181083 |
| Mouse monoclonal anti-GAPDH (loading control, clone "GAPDH 1D4") | Imgenex (India) | Cat# IMG-5019A-1, RRID:AB_316884 |
| Mouse monoclonal anti-DUX4 (clone P4H2) | Novus Biologicals (CO, USA) | Cat# NBP1-49552 |
| Rabbit polyclonal anti-UPF1 | Cell Signaling Technology (MA, USA) | Cat# 9435, RRID:AB_10629662 |
| Rabbit recombinant monoclonal anti-eIF4A3 (clone EPR14301(B)) | Abcam (MA, USA) | Cat# ab180573, RRID:AB_370675 |
| Rabbit polyclonal anti-CWC22 | Novus Biologicals via Antibody Registry | Cat# H00057703-B01P, RRID:AB_2087011 |
| Rabbit polyclonal anti-PRP8 (PRPF8) | Abcam (MA, USA) | Cat# ab79237 |
| Rabbit polyclonal anti-UPF3B (UPF3B/UPF3A) | Abcam (MA, USA) | Cat# ab269998, |
| Rabbit IgG (H+L) control antibody | Rockland Immunochemicals (USA) | Cat# ab46540, RRID:AB_2614925 |
